# Tumor infiltrating iNKT cells sustain neutrophil pro-tumorigenic functions influencing disease progression in human colorectal cancer

**DOI:** 10.1101/2022.10.19.512854

**Authors:** Georgia Lattanzi, Francesco Strati, Angélica Díaz-Basabe, Federica Perillo, Chiara Amoroso, Giulia Protti, Maria Rita Giuffrè, Ludovica Baldari, Elisa Cassinotti, Michele Ghidini, Barbara Galassi, Gianluca Lopez, Daniele Noviello, Laura Porretti, Elena Trombetta, Luca Mazzarella, Giandomenica Iezzi, Francesco Nicassio, Francesca Granucci, Flavio Caprioli, Federica Facciotti

**Affiliations:** Department of Experimental Oncology, European Institute of Oncology IRCCS, Milan, Italy; Department of Oncology and Hemato-oncology, Università degli Studi di Milano, Milan, Italy; Department of Biotechnology and Biosciences, University of Milano-Bicocca, Milan, Italy; Gastroenterology and Endoscopy Unit, Fondazione IRCCS Cà Granda, Ospedale Maggiore Policlinico, Milan, Italy; General and Emergency Surgery Unit, Fondazione IRCCS Ca’ Granda, Ospedale Maggiore Policlinico, Milan, Italy; Medical Oncology, Fondazione IRCCS Ca’ Granda, Ospedale Maggiore Policlinico, Milan, Italy; Pathology Unit, Fondazione IRCCS Cà Granda, Ospedale Maggiore Policlinico, Milan, Italy; Department of Pathophysiology and Transplantation, Università degli Studi di Milano, Milan, Italy; Clinical Chemistry and Microbiology Laboratory, Fondazione IRCCS Ca’ Granda Ospedale Maggiore Policlinico, Milan, Italy; Department of Visceral Surgery, EOC Translational Research Laboratory, Bellinzona, Switzerland; Center for Genomic Science of IIT@SEMM, Istituto Italiano di Tecnologia (IIT), Milan, Italy

**Keywords:** iNKT, CRC, neutrophils, *Fusobacterium nucleatum*

## Abstract

iNKT cells account for a relevant fraction of effector T-cells in the intestine. Although iNKT cells are cytotoxic lymphocytes, their role in colorectal cancer (CRC) remains controversial. From the analysis of colonic LPMCs of human and murine CRC specimens we report that tumor-infiltrating iNKT cells are characterized by an IL17/GM-CSF pro-tumorigenic phenotype, while maintaining cytotoxic properties in the adjacent non-tumoral tissue. Exposure of iNKT cells to the tumor-associated pathobiont *Fusobacterium nucleatum* blunted their cytotoxic capability and enhanced iNKT cell-mediated neutrophils chemotaxis, which upregulated PMN-MDSC gene signatures and functions. Importantly, *in vivo* stimulation of iNKT cells with αGalCer restored their anti-tumorigenic functions. Survival analyses demonstrated that human CRC co-infiltration by iNKT cells and tumor-associated neutrophils correlates with negative outcomes.

Our results reveal a functional plasticity of human intestinal iNKT cells with pro- and anti-tumorigenic activities in CRC, suggesting an iNKT pivotal role in shaping the cancer developmental trajectory.

## Introduction

Invariant Natural Killer T cells (iNKT) are a lipid-specific, evolutionary conserved, population of lymphocyte positioned at the interface between innate and adaptive immunity ^1,2^. Microbial and endogenous ^3,4^ signals finely tune iNKT cell functions, including tissue immune surveillance ^5^ and first line defense against infectious microorganisms ^1^. iNKT cells are present also in the intestinal lamina propria as tissue resident cells and variations in the gut microbiota composition can rapidly alter their phenotype ^1,6^. Dysbiosis imprints iNKT cells toward a pro-inflammatory phenotype ^7^ whereas normobiosis restoration upon fecal microbiota transplantation ^8,9^ and exposure to Short Chain Fatty Acids (SCFA) ^8,9^ induce their production of regulatory cytokines, such as IL10. Along with the patrolling of tissue integrity, iNKT cells actively participate in the immune surveillance against malignant transformation and tumor progression, including human colorectal cancer (CRC) ^10^. CRC is the third most prevalent cancer worldwide and the second leading cause of cancer-related death ^11^. Microbiota-elicited inflammation is an important contributor to CRC pathogenesis regardless of pre-cancer inflammatory history ^12^. Because of their fast responsiveness to microbes, iNKT cells can produce a large amount of effector cytokines ^1^ during the time required for the recruitment, activation and expansion of conventional T cells ^13^ with the potential to robustly imprint the tumor microenvironment (TME) and the CRC developmental trajectory. iNKT cells are considered important in antitumor immunity ^10^ and their infiltration in tumoral lesions is a positive prognostic factor in different cancer types ^14,15^. Moreover, iNKT cells possess cytotoxic properties ^16^ and are endowed with cell-killing activities towards different human CRC cell lines and primary patient’s derived cancer epithelial cells through the perforin– granzyme pathway ^17^. However, their role in CRC progression has never been fully elucidated and it is still controversial ^14,18^. Indeed, iNKT cells with a pro-tumorigenic phenotype have been described in murine models of CRC and associated with shorter disease-free survival in patients ^18,19^. Here, by taking advantage of a large cohort of human CRC specimens and of different murine models of colon carcinogenesis, we addressed the contribution of iNKT cells to CRC pathophysiology and the effects of tumor-associated microbiota in shaping their cytotoxic potential. We demonstrate that tumor-infiltrating iNKT cells, but not those isolated form adjacent tumor-free areas, manifest a pro-tumorigenic phenotype and correlate with negative disease outcomes in CRC patients. We show that the CRC-associated pathobiont *F. nucleatum* blunts the cytotoxic functions of iNKT cells leading to the recruitment in the TME of neutrophils with phenotypic and functional characteristics ascribable to polymorphonuclear myeloid-derived suppressor cells (PMN-MDSCs). Finally, we show that restoring the cytotoxic potential of iNKT cells by *in vivo* treatment with the iNKT-specific agonist α-galactosylceramide (αGalCer) leads to control of tumor growth.

## Results

### Tumor-infiltrating human iNKT cells show a pro-tumorigenic profile and correlate with TANs infiltration

To uncover the role of iNKT cells in human CRC, we collected freshly isolated surgical specimens from 118 CRC patients enrolled at the Policlinico Hospital Milan, whose clinical features are described in Table 1.

**Table 1:**
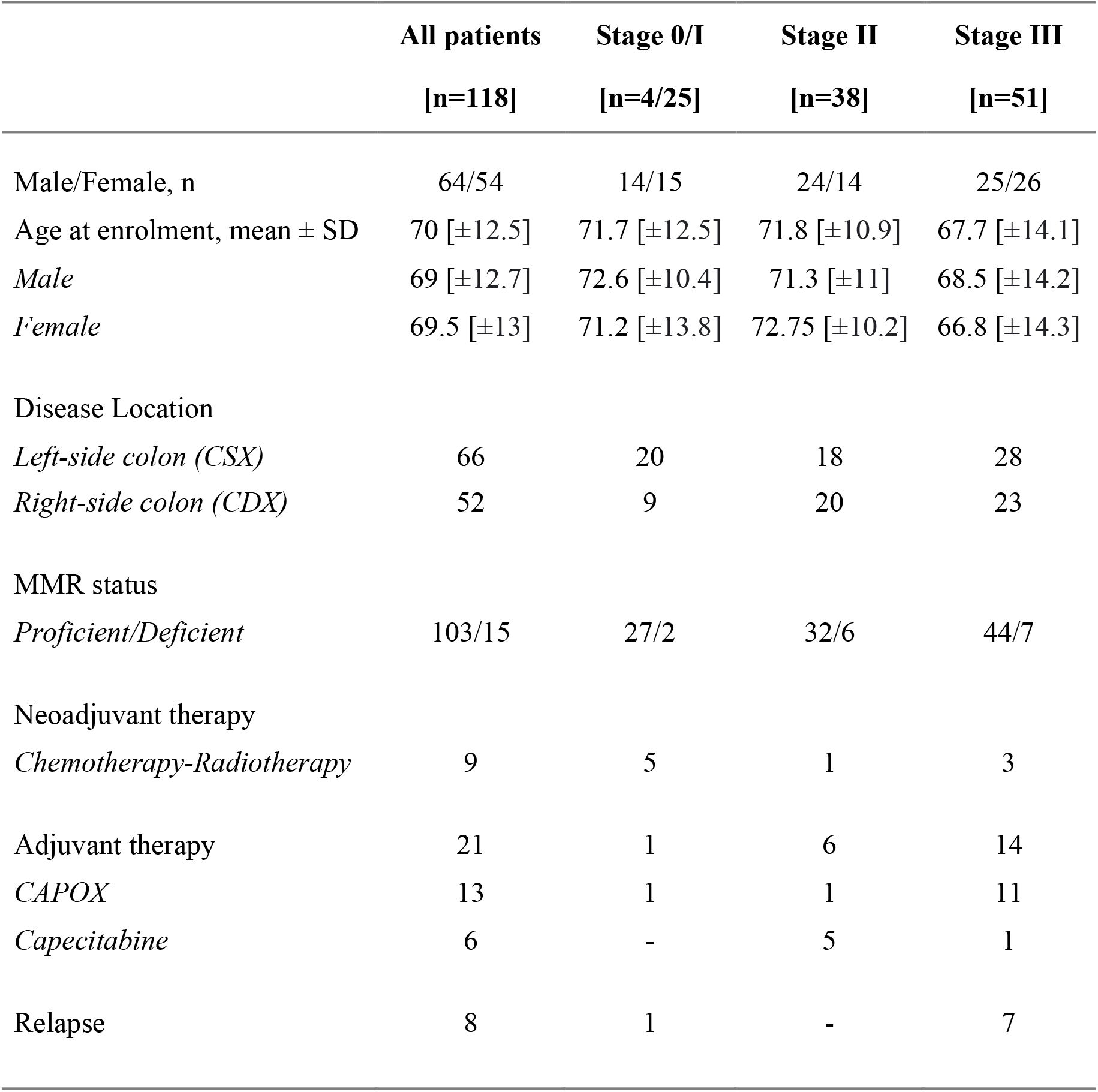
Clinical characteristics of the study population.

Multidimensional immunophenotyping of tumor lesions (TUM) as well as adjacent non-tumor colon tissue (NCT) revealed that iNKT cells are significantly enriched in TUM samples (Figure 1A, Supplementary Figure 1A). By applying a Phenograph unsupervised clustering (Supplementary Figure 2A-B), we identified a cluster of iNKT cells (cluster C16) specifically enriched in TUM (Figure 1B, Supplementary Figure 2A-B); the metaclustering analysis of cluster C16 revealed that tumor-infiltrating iNKT cells are characterized by an overall increased expression of IL17 and GM-CSF (clusters C1 and C2) whereas IFNγ is expressed mainly by NCT-infiltrating iNKT cells (cluster C4, Figure 1C). Manual gating analysis confirmed the increased frequency of GM-CSF^+^IL17^+^iNKT cells in CRC lesions and of IFNγ^+^iNKT cells in NCT (Figure 1D-E). The co-expression of IL17 and GM-CSF was a unique feature of iNKT cells, distinguishing them from other tumor-infiltrating conventional (CD4^+^ and CD8^+^) and unconventional (γδ T and MAIT cells) T cells (Supplementary Figure 2C-D). Phenotypically, tumor-infiltrating iNKT cells displayed an increased expression of exhaustion/inhibitory molecules including PD-1, TIGIT and TIM-3 (Supplementary Figure 2E) and a reduced expression of activation markers such as CD69, CD161 and CD137 (4-1BB) as compared to NCT (Supplementary Figure 2F). No differences were observed for the secretion of other cytokines and cytotoxic molecules compared to NCT, although we observed a decreased expression of the pro-apoptotic factor Fas ligand (FasL) in tumor-infiltrating iNKT cells (Supplementary Figure 2G-H). These data collectively indicate that human tumor-infiltrating iNKT cells are skewed towards a pro-tumorigenic phenotype, characterized by the secretion of GM-CSF, IL17 and the expression of exhaustion/inhibitory molecules.

**Figure 1:**
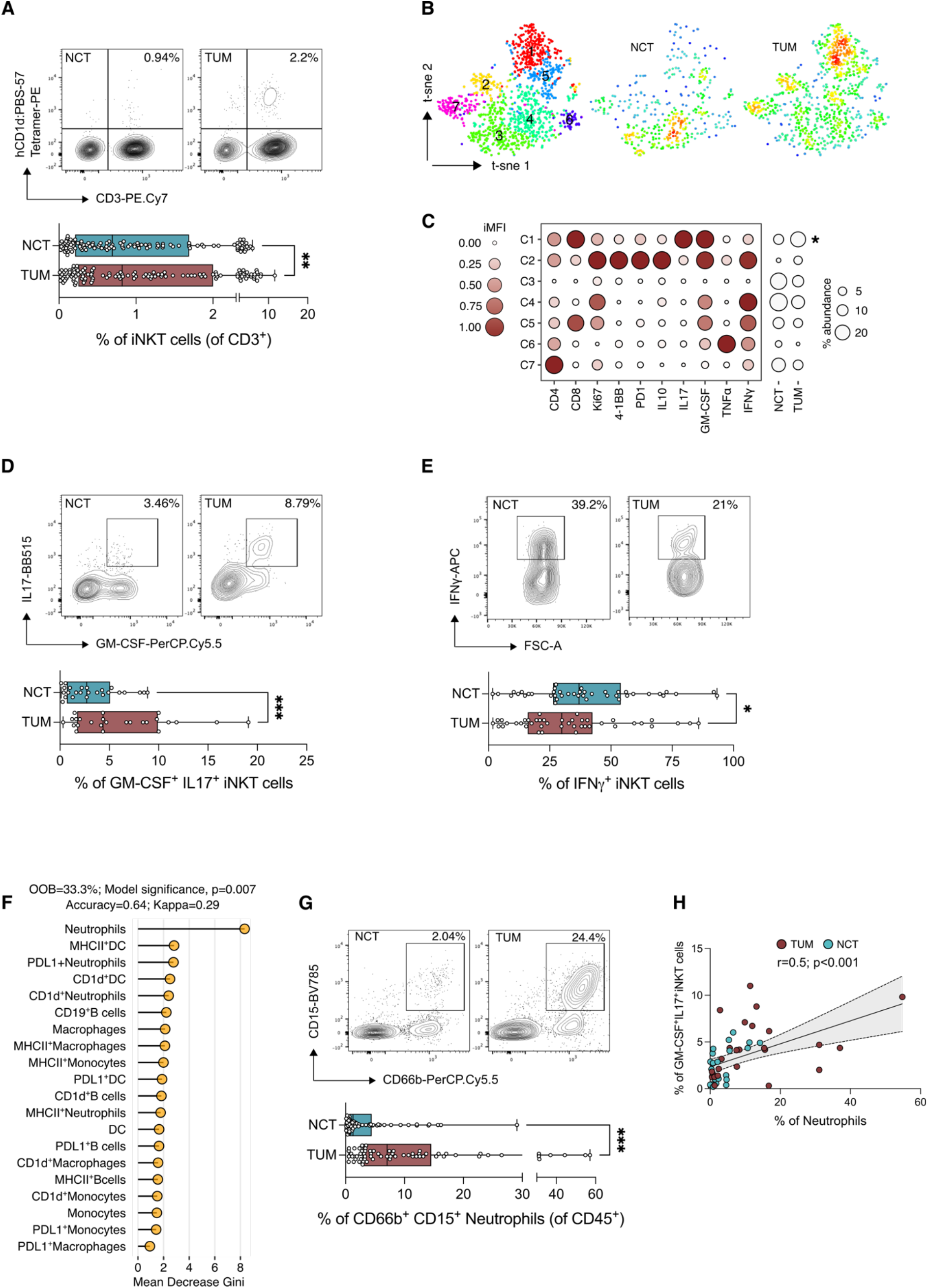
iNKT cells infiltrate CRC lesions and correlate with neutrophil abundance. **(A)** Frequency of iNKT cells in adjacent non-tumor colonic tissue (NCT, blue boxes) and tumor lesions (TUM, red boxes) (n=115), with representative dot plots. **(B)** t-SNE map of iNKT cells based on Phenograph metaclustering analysis in NCT and TUM samples. **(C)** Balloon plot of the scaled integrated Mean Fluorescent Intensity (MFI) of Phenograph clusters generated in B. (**D, E)** Frequency of IL17^+^GM-CSF^+^ **(D)** and IFNγ^+^ **(E)** iNKT cells infiltrating NCT and TUM. **(F)** Random Forest analysis of myeloid and B cell compartment in NCT and TUM with the highest discriminatory power, sorted by mean decrease GINI value. **(G)** Frequency of neutrophils in NCT and TUM (n=75) with representative plots. (**H**) Spearman’s correlation analysis of IL17^+^GM-CSF^+^iNKT and neutrophils infiltrating NCT and TUM (n=25). P < 0.05 (*), P < 0.01 (**), P < 0.001(***); Wilcoxon signed-rank test.

iNKT cells are able to modulate the activity of myeloid cells during homeostasis, inflammation and tumor development ^20^. Thus, we hypothesized that tumor-infiltrating iNKT cells may shape the TME by acting primarily on innate immune cells. A random forest-based classification modelling identified neutrophils (CD45^+^CD66b^+^CD15^+^ cells, Supplementary Figure 1B) as the most important innate immune cell population (Supplementary Figure 3A) to classify samples according to their location (*i.e*., TUM vs NCT) (Figure 1F). Neutrophils were significantly enriched in CRC lesions (Figure 1G), had a mature (CD33^mid^CD10^+^CD16^+^), aged-like phenotype (CD62L^-^CXCR4^+^) and downregulated the expression of the antigen presenting molecules CD1d and MHC-II (Supplementary Figure 3B-C). No differences were detected in the expression of the immune checkpoint Programmed Death-Ligand 1 (PD-L1, Supplementary Figure 3C). Most importantly, neutrophils correlated with the frequency of GM-CSF^+^IL17^+^iNKT cells (Spearman’s r=0.5, p<0.001, Figure 1H), but not with total iNKT cells (Supplementary Figure 3D) suggesting a specific crosstalk between Th17-like iNKT cells and tumor-associated neutrophils (TANs). Altogether these data suggest the existence of a functional iNKT-TAN axis in CRC.

### Tumor-associated *Fusobacterium nucleatum* induces a pro-tumorigenic signature in iNKT cells and favors neutrophil recruitment

The gut microbiota is an oncogenic driver of CRC ^21^ and intestinal microbes represent potent stimulators of iNKT cell responses ^7^. Thus, we analyzed the tumor-associated microbiota by 16S rRNA gene sequencing and identified *Fusobacterium* as one of the most enriched Amplicon Sequence Variant (ASV) in TUM vs NCT (Figure 2A). *Fusobacterium nucleatum* (*Fn*) is a hallmark of CRC, extensively studied for its pro-tumorigenic properties ^21^, but its effect on iNKT cells has never been tested. Thus, we primed human iNKT cell lines with monocyte-derived dendritic cells (moDC) pulsed with *Fn* or αGalCer, the prototype agonist of iNKT cells ^22^, and performed *in vitro* functional and cytotoxic assays as well as RNA sequencing of iNKT cells (Figure 2B). Exposure of iNKT cells to *Fn* impaired their *in vitro* cytotoxic functions against colon adenocarcinoma cell lines (Figure 2C). *Fn*-primed iNKT cells showed an enriched neutrophil chemotaxis gene signature, which included the chemokines of the C-X-C and C-C motif ligand family genes *CXCL8, CXCL2, CXCL3, CCL3L1, CCL4L2, CCL20* and *CCL22* (Figure 2D-E). By contrast, αGalCer-primed iNKT cells presented an IFNγ/cytotoxic-related gene signature, including the expression of *TBX21, IFNG, PFN1, GNLY, GZMA, GZMB, GZMH, LTA, LTB* and *NKG7* (Figure 2D). Consistently αGalCer-primed iNKT cells secreted IFNγ while *Fn* induced the production of GM-CSF and IL17 (Figure 2F).

**Figure 2:**
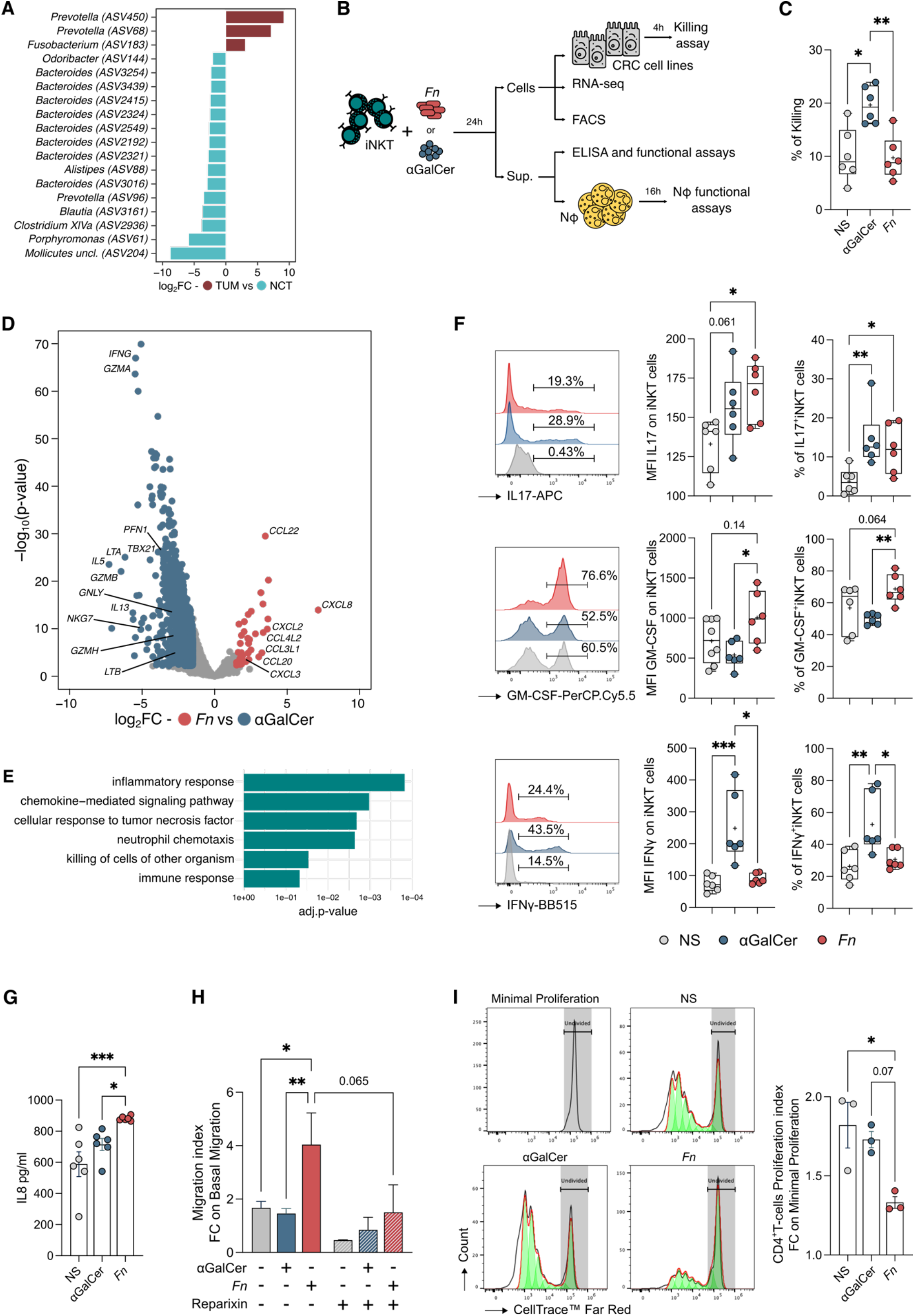
*Fusobacterium nucleatum* reduces iNKT cell cytotoxicity and promotes iNKT-cell dependent activation of neutrophils. **(A)** Bar plot representing the significantly enriched amplicon sequence variants (ASVs) (FDR p <0.05) in TUM vs NCT mucosal samples (n=70) by DEseq2 analysis. **(B)** Schematic representation of the experimental plan. **(C)** Percentage of killing of unstimulated (NS), αGalCer- or *F. nucleatum* (*Fn*)-primed iNKT cells; results are representative of three (n=3) independent experiments **(D)** Volcano plot representing the differentially expressed genes (DEGs) in *Fn*- vs αGalCer-primed iNKT cells; the volcano plot shows for each gene (dots) the differential expression (log_2_fold-change [log_2_FC]) and its associated statistical significance (log_10_p-value). Dots indicate those genes with an FDR-corrected p < 0.05 and log2FC > |1.5|. **(E)** Gene Ontology (GO) analysis of the differentially expressed genes (Bonferroni-corrected p < 0.05 and log_2_FC > 1). **(F)** Representative histograms (left panels), MFI (middle panels) and frequency (right panels) of IL17^+^, GM-CSF^+^ and IFNγ^+^ iNKT cells unstimulated (NS) or primed with either αGalCer or *Fn*. **(G)** hIL8 protein concentration in the supernatant released by unstimulated (NS) or αGalCer- or *Fn*-primed iNKT cells. (**H)** Fold-change of neutrophils migration index upon exposure to unstimulated (gray bar), αGalCer (blue bars) or *Fn*-primed (red bars) iNKT cell supernatants in the absence (full bars) or presence (pattern fill bars) of Reparixin (20 μM). (**I**) Proliferation index of näive CD4^+^T cells co-cultured with neutrophils and cell free supernatants from unstimulated (NS), αGalCer or *Fn* primed-iNKT cells. P < 0.05 (*), P < 0.01 (**), P < 0.001(***); Kruskal-Wallis test. Data are representative of at least three independent experiments.

Since iNKT cells may impact on neutrophil survival, recruitment and activation status ^23–25^, we evaluated how *Fn* affected the crosstalk between iNKT cells and neutrophils. Both *Fn*- and αGalCer-primed iNKT cells increased the survival rate of neutrophils compared to unstimulated cells (Supplementary Figure 4A). However, only *Fn*-primed iNKT cells induced neutrophil recruitment (Figure 2H), in line with the upregulation of *CXCL8* (Figure 2D) and the higher concentration of IL8 in the iNKT cell-derived culture supernatant (Figure 2G). Neutrophil migration was inhibited by the use of Reparixin, *i.e*., an allosteric inhibitor of the IL8 receptor (CXCR-1/-2), demonstrating that neutrophil chemotaxis is affected by chemokines produced by *Fn*-primed iNKT cells (Figure 2H). Moreover, *Fn*-primed iNKT cells affected neutrophil activation status by reducing their respiratory burst capability, inducing the expression of PD-L1 (Supplementary Figure 4C-D) and promoting their suppressive activity towards CD4^+^T cell proliferation (Figure 2I). The above analyses suggest that iNKT cells conditioning by CRC-associated microbiota promotes recruitment and induces an immunosuppressive phenotype in neutrophils.

### The absence of iNKT cells limits tumor burden by reducing pro-tumorigenic TANs

To functionally dissect the dynamic interaction between iNKT cells and neutrophils in CRC we used two different CRC murine models: the chemical azoxymethane-dextran sodium sulphate (AOM-DSS) model of colitis-associated CRC and the syngeneic MC38 model.

Since mucosal iNKT cells are largely tissue-resident lymphocytes ^13^, they might infiltrate tumors at the beginning of their formation and, given their functional interaction with TANs, affect the TME by modulating neutrophil behavior. To test this hypothesis, we first evaluated the dynamics of tumor growth and intratumor frequency of iNKT cells and neutrophils in the AOM-DSS model (Supplementary Figure 5A-B). Tumor-infiltrating iNKT cells reached their peak abundance between day 21 and day 28 from tumor induction (Supplementary Figure 5A). Conversely, the kinetic of neutrophils infiltration started at day 35 and increased until day 42; then, both neutrophils and iNKT cells abundance declined (Supplementary Figure 5A). At this timepoint, *i.e*., day 49 from tumor induction (T1), the AOM-DSS model mirrored the key phenotypic and functional features of tumor infiltrating iNKT cells (Supplementary Figure 5C-D) and neutrophils (Supplementary Figure 5E-F) observed in our human cohort. Interestingly, at later timepoints from tumor induction *i.e*., day 70 (T2), tumor-infiltrating iNKT cells began to lose the key features observed in CRC patients (Supplementary Figure 5G-H) prompting us to focus on earlier timepoints *in vivo*.

To rule out the anti- or pro-tumorigenic role of iNKT cells in CRC, we induced tumorigenesis in animals lacking iNKT cells *i.e*., *CD1d*^-/-^ and *Traj18*^-/-^ mice. Both iNKT cell-deficient strains showed a reduced tumor formation compared to wild-type C57BL/6 (B6) mice (Figure 3A-C). The abundance of TANs was significantly reduced in *CD1d*^-/-^ and *Traj18*^-/-^ compared to iNKT-cell proficient mice (Figure 3D). Moreover, *Traj18*^-/-^ TANs negatively correlated with the number of tumors whereas tumors increased proportionally with TANs in B6 mice (Figure 3E). Next, we sought to understand if iNKT cells could shape distinct biological programs and unique molecular features of TANs. Transcriptomics analysis of sorted CD45^+^Lin^-^ CD11b^+^Ly6G^+^ cells from B6 and *Traj18*^-/-^ mice revealed that TANs from B6 animals were enriched for transcripts of chemokines and inflammation-related molecules (*Ccl3, Cxcl2, Cxcr5, Nfkbie, Nfkbiz, Socs3, Atf4, Ptsg2, Pla2g7*) as well as of immune suppression (*Il10, S100a8*) (Figure 3F and 3H); these genes have all been associated with different populations of polymorphonuclear myeloid-derived suppressor cells (PMN-MDSCs) ^26,27^. Additional MDSC markers identified include the C-type lectin domain family 4-member N and D (*Clec4n* and *Clec4d*), activating protein 1 transcription factor subunit (*Junb*) and myeloid cell surface antigen CD33 (*Cd33*) ^27^. On the other hand, *Traj18*^-/-^ -isolated TANs showed a marked increased expression of genes associated with MAPK signaling (*Map3k14, Map14, Map12*), NETs release (*Hmgb1, Hmgb2, Ceacam1, Mmp15, Mmp21*) ^28,29^, the hypoxia inducible factor 1 subunit alpha (*Hif1a*) and the anti-apoptotic BAG cochaperone 4 (*Bag4*) (Figure 3F and 3H). Pathway enrichment analysis highlighted the upregulation of genes associated with TNF signaling, mostly in B6 TANs (Figure 3G). To understand if the differential transcriptional activity of TANs in B6 and *Traj18*^-/-^ mice could be restricted to different populations of PMN-MDSCs, we analyzed a publicly available scRNA-seq dataset of PMN-MDSC from tumor-bearing mice ^26^. The t-SNE overlay analysis revealed the enrichment of the B6 TANs gene signature in a cluster of ‘activated’ PMN-MDSCs (PMN3) while the *Traj18*^-/-^ TANs gene signature was associated with a population of immature neutrophils/classical PMN-MDSCs (PMN2), reflecting different pathways of PMN-MDSC activation ^26^ in the presence or absence of iNKT cells (Supplementary Figure 6). Accordingly, we identified two distinct and differentially represented populations of TANs in B6 and iNKT cell-deficient animals. *CD1d*^-/-^ and *Traj18*^-/-^ mice showed a lower frequency of CD11b^+^Ly6g^low^TANs and an increase of CD11b^+^Ly6g^high^TANs (Figure 3I). CD11b^+^Ly6g^low^ cells had reduced respiratory burst capacity (Figure 3J) suggesting a diminished cytotoxic potential ^30^, and increased expression of PD-L1 compared to CD11b^+^Ly6g^high^ cells (Figure 3K). Collectively, these findings reveal that intestinal iNKT cells are associated with different functional subgroups of neutrophils *in vivo*, that may play a role in CRC progression.

**Figure 3:**
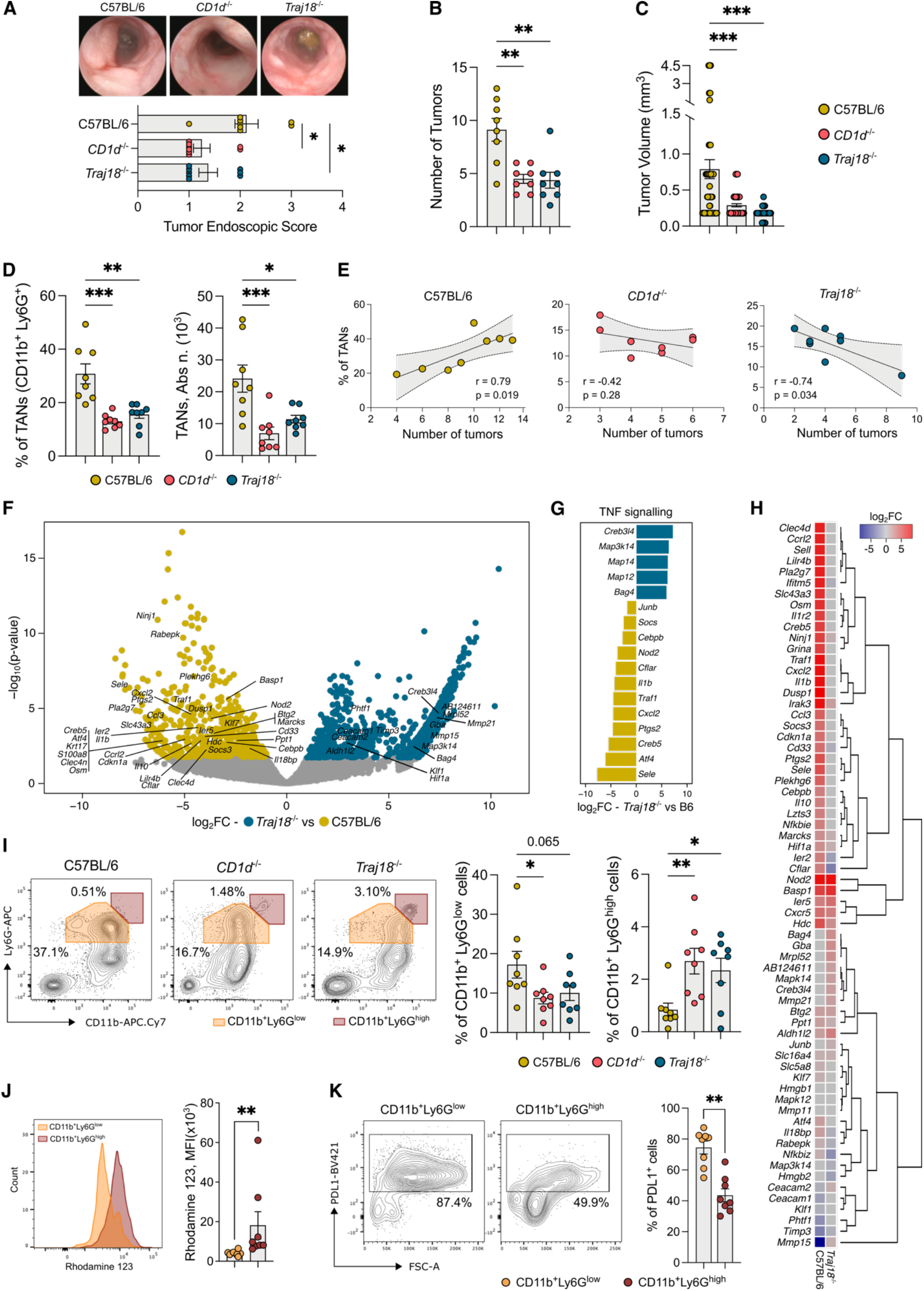
The absence of iNKT cells decreases tumor formation *in vivo* and associates with a less pro-tumorigenic gene signature in TANs. **(A-C)** Cumulative tumor endoscopic score and representative endoscopic pictures **(A)**, number **(B)** and volume **(C**) of tumors from AOM-DSS treated C57BL/6, *CD1d*^-/-^ and *Traj18*^-/-^ animals **(D)** Frequencies (left panels) and absolute numbers (right panels) of TANs in C57BL/6, *CD1d*^-/-^ and *Traj18*^-/-^ animals. **(E)** Correlation analysis of TANs frequency and number of tumors in C57BL/6, *CD1d*^-/-^ and *Traj18*^-/-^ animals. **(F)** Volcano plot representing the DEGs of TANs in C57BL/6 and *Traj18*^-/-^ animals; the volcano plot shows for each gene (dots) the differential expression (log_2_fold-change [log_2_FC]) and its associated statistical significance (log_10_p-value). Dots indicate those genes with an FDR-corrected p < 0.1 and log_2_FC > |1|. **(G)** DEGs enriched in the KEGG TNF signaling pathway (Bonferroni-corrected p < 0.05 and log_2_FC > |1|). **(H)** Heatmap and hierarchical clustering of MDSC-related DEGs (FDR-corrected p-value < 0.05 and log_2_FC > |1|) in neutrophils from C57BL/6 and *Traj18*^-/-^ tumor bearing vs healthy controls **(I)** Frequency of CD11b^+^, Ly6G^high^ and Ly6G^low^ TANs in C57BL/6, *CD1d*^-/-^ and *Traj18*^-/-^ animals, with representative dot plots. **(J, K)** Respiratory burst quantification **(J)** and frequency of PD-L1^+^ **(K)** in CD11b^+^Ly6G^high^ and CD11b^+^Ly6G^low^ TANs in *Traj18*^-/-^ mice, with representative plots. Data points (n=8) from two pooled independent experiments representative of at least three. P < 0.05 (*), P < 0.01 (**), P < 0.001(***); Kruskal-Wallis and Mann-Whitney tests. Two-tailed Pearson test for correlation analysis.

### *In vivo* αGalCer treatment reactivates the cytotoxic potential of iNKT cells

Next, we asked whether restoring the activation status of iNKT cells towards a cytotoxic phenotype could remodel the cancer developmental trajectory in the MC38 CRC model (Figure 4A). First, we confirmed the pro-tumorigenic role of iNKT cells, since *Traj18*^-/-^ mice showed a significantly delayed tumor growth compared to B6 animals (Figure 4B-D) and a reduced infiltration of TANs (Figure 4E). Then, we tested whether the *in vivo* administration of αGalCer to MC38-bearing B6 mice could revert the pro-tumorigenic phenotype of iNKT cells. We found that αGalCer treatment significantly reduced tumor growth (Figure 4F-H) re-establishing iNKT cells ability to express high levels of IFNγ (Figure 4I); no difference was observed for GM-CSF and IL17 secretion in treated and untreated MC38-bearing mice (Figure 4J) suggesting that an intestinal microenvironment may be relevant to polarize this phenotype in iNKT cells. Indeed, *in vitro* priming of splenic iNKT cells with the gut microbiota of AOM-DSS treated mice (B6, *CD1d*^-/-^ and *Traj18^-/-^*) induced IL17 and GM-CSF expression on iNKT cells (Supplementary Figure 7A). Notably, the gut microbiota of healthy mice had no effect on IL17 and GM-CSF production (Supplementary Figure 7A) suggesting that only a dysbiotic, CRC-associated microbiota (Supplementary Figure 7B-C) can promote a GM-CSF^+^IL17^+^iNKT cell phenotype.

**Figure 4:**
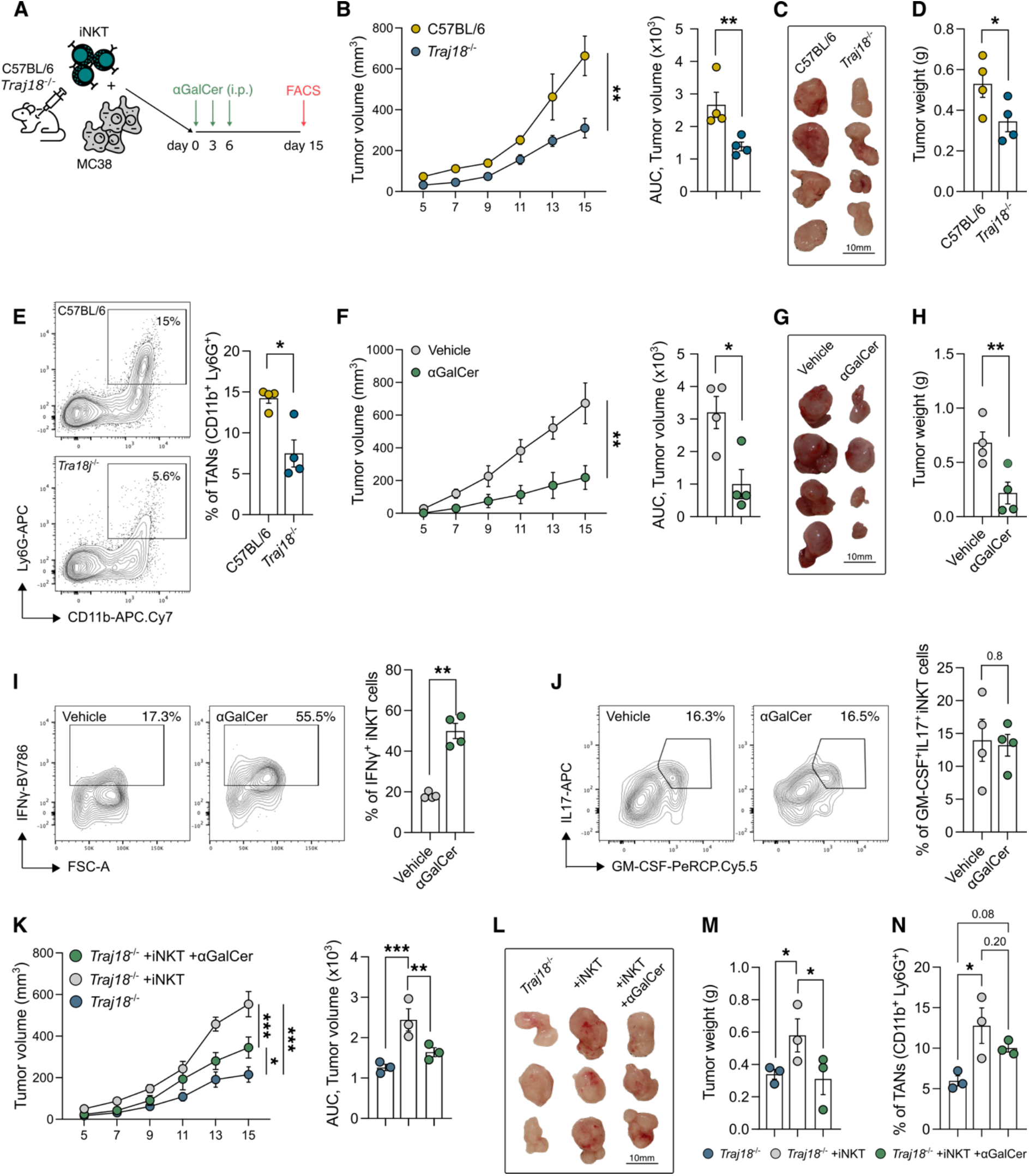
*In vivo* αGalCer administration restores iNKT cell anti-tumorigenic functions. **(A)** Schematic representation of experimental plan. **(B)** MC38 tumor growth in C57BL/6 and *Traj18*^-/-^ animals, with their relative area under the curve (AUC). **(C-D)** MC38 tumors representative pictures **(C)** and weight of tumors **(D)** from MC38-bearing C57BL/6 and *Traj18*^-/-^ animals. **(E)** Frequency of TANs in MC38-bearing C57BL/6 and *Traj18*^-/-^ animals, with representative dot plots. **(F-H)** MC38 tumor growth with their relative AUC **(F)**, representative pictures **(G)** and weight of tumors **(H)** from MC38-bearing C57BL/6 mice treated, or not, with αGalCer. **(I, J)** Frequency of tumor infiltrating IFNγ^+^ **(I)** and GM-CSF^+^IL17^+^ **(J)** iNKT cells in MC38-bearing C57BL/6 animals treated, or not, with αGalCer, with representative dot plots. **(K-N)**, MC38 tumor growth and AUC (**K)**, representative pictures **(L)**, weight of tumors (**M)** and frequency of TANs **(N)** in MC38-bearing *Traj18*^-/-^ animals reconstituted, or not, with iNKT cells prior to treatment with αGalCer. Data shown (n=3-4) are representative of at least one of three independent experiments. P < 0.05 (*), P < 0.01 (**), P < 0.001(***). Kruskal-Wallis test and Two-Way ANOVA for tumor growth.

Finally, reconstitution of *Traj18*^-/-^ mice with splenic iNKT cells promoted tumor growth and TANs infiltration (Figure 4K-N), although αGalCer-mediated iNKT cell activation effectively reverted the pro-tumorigenic effects of iNKT cells on tumor growth (Figure 4K-M). These data confirm that the deleterious effects exerted by the TME on iNKT cells, and consequently their support to tumorigenesis, can be reverted by manipulating their activation status.

### iNKT cell infiltration impairs the favorable prognostic significance of TANs in human CRC and correlates with poor patient outcomes

Our findings identified a pro-tumorigenic role for iNKT cells in CRC and a cross-talk with TANs in murine models. To contextualize the significance of these results in human CRC, we stratified our patient cohort in tumor-infiltrating iNKT^high^ and iNKT^low^ subgroups and performed Kaplan-Meier analyses. Relapse free survival (RFS) at 4 years was higher in iNKT^low^ CRC patients (Figure 5A). Several studies described a favorable prognostic significance for neutrophil infiltration in CRC ^31–33^, which we confirmed in our cohort (Figure 5B); however, the neutrophil positive prognostic significance in CRC was lost in iNKT^high^ patients (Figure 5C) thus indicating that the beneficial effects of neutrophils on clinical outcomes require the concomitant low infiltration of iNKT cells. Moreover, we validated these results by interrogating the colon adenocarcinoma cohort (COAD) of The Cancer Genome Atlas (TCGA) database ^34^ and found that the positive prognosis associated with neutrophil infiltration in CRC patients (as measured by the expression of the *CEACAM8* gene, encoding for CD66b) was dependent on the low expression of the iNKT cell specific transcription factor PLZF, encoded by the *ZBTB16* gene ^35^ (Figure 5D-E).

**Figure 5:**
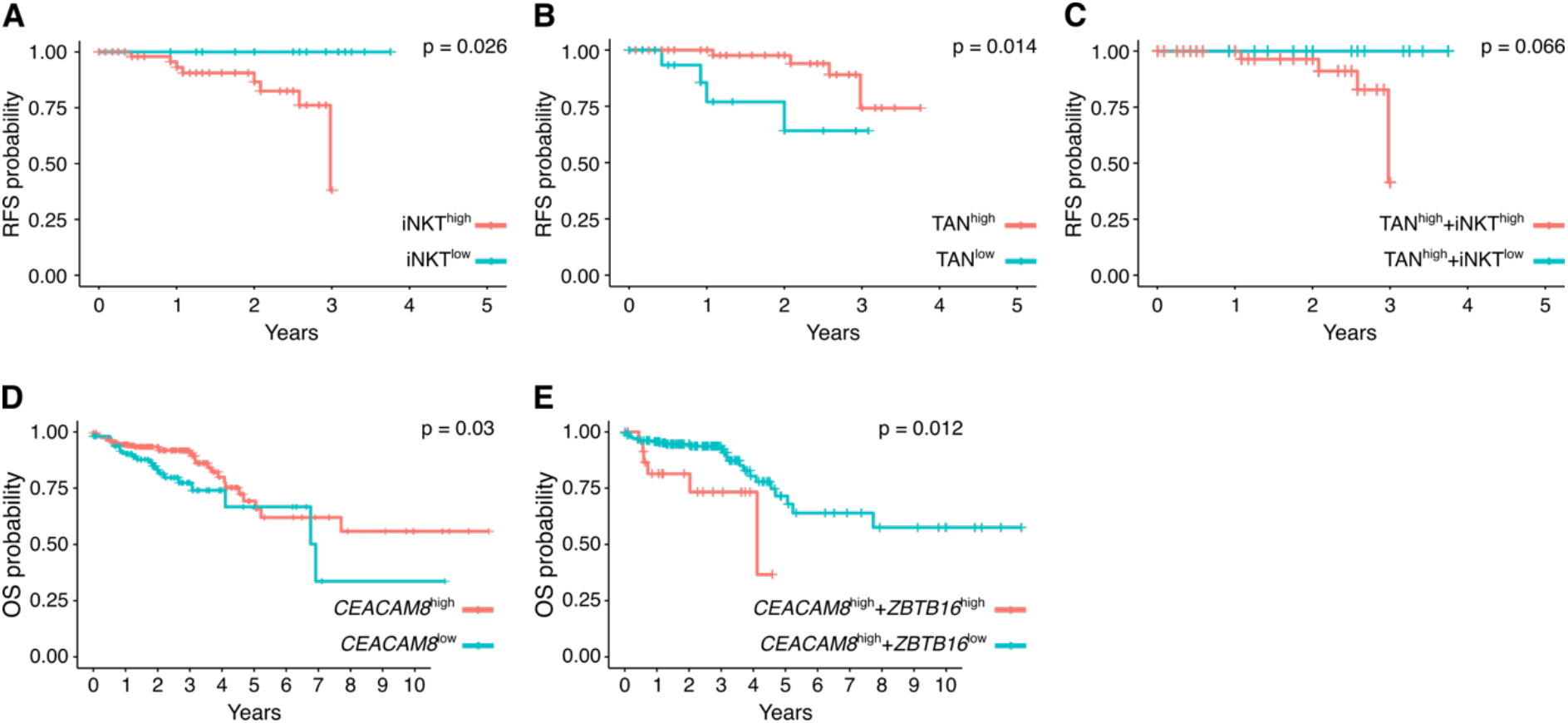
iNKT cell infiltration correlates with poor patient outcomes. **(A-C)** Kaplan-Meier relapse free survival (RFS) curves of CRC patients from Policlinico Hospital, Milan presenting high vs low (**A)** tumor infiltrating iNKT cells, **(B)** high vs low TANs or **(C)** high vs low tumor infiltrating iNKT cells in the population of TAN^high^ patients. **(D-E)** Kaplan-Meier overall survival (OS) curves of CRC patients from TCGA cohort with respect to **(D)** high or low expression of *CEACAM8* within tumor specimens and **(E)** high or low expression of *ZBTB16* in the in the population of *CEACAM8*^high^ patients. **(F)** Proposed model for iNKT-mediated pro/antitumor immunity in CRC.

## Discussion

iNKT cells are essential components of anti-tumor responses thanks to their well-known cytotoxic properties ^16^ and active participation in immune surveillance against malignant transformation ^10^. However, reduced frequencies and functional impairment of iNKT cells have been associated with poor overall survival in solid or hematological tumors ^10^ associated with loss of IFNγ secretion and acquisition of an anti-inflammatory phenotype ^36^. Nevertheless, αGalCer administration can revert iNKT cells functional impairment ^37^ as demonstrated in preclinical studies and RCTs ^38^. Intriguingly, iNKT cells showed also pro-tumorigenic functions, as reported in the spontaneous murine adenomatous polyposis coli *Apc^Min/+^* model for colon cancer, where iNKT cells promote tumor progression ^19^.

Here, we further expanded these findings by analyzing human CRC specimens and different murine models and describing the opposing roles of human and murine iNKT cells in paired non-tumor vs cancerous tissue. In this study, the immunophenotypic profiles of *ex vivo* isolated human iNKT cells and the use of two different iNKT cell-deficient murine strains confirmed that IFNγ-producing, cytotoxic iNKT cells limit colonic tumorigenesis whereas intratumoral accumulation of GM-CSF^+^IL17^+^iNKT cells support colon cancer progression.

The gut microbiota is an oncogenic driver of CRC ^21^ and intestinal microbes represent potent stimulators of iNKT cell responses that can shape their functional plasticity ^7^. *Fusobacterium* is a hallmark CRC-associated pathobiont ^21^ we found enriched in our patients’ cohort. We showed that iNKT cell stimulation with *F. nucleatum* promotes their commitment towards the production of IL17 and GM-CSF impairing iNKT cell cytotoxicity. *F. nucleatum* is known to suppress anti-tumor immunity by binding to the TIGIT receptor on NK cells through the virulence factor Fap2 ^39^. Accordingly, we observed increased TIGIT expression in tumor-infiltrating iNKT cells. Moreover, *F. nucleatum* imprinted a neutrophil chemotaxis gene signature in iNKT cells, with *CXCL8* being the most upregulated gene. CXCL8, namely IL8, is a chemokine that promotes neutrophil migration which is expressed also by TILs ^40^. By expressing CXCL8, iNKT cells may regulate neutrophils trafficking within the TME, shaping early immune responses in CRC. Our *in vitro* observations and the early tumor infiltration of iNKT cells with respect to neutrophils *in vivo* suggest that this hypothesis may be valid. Few studies reported the close interaction of iNKT cells with neutrophils. In melanoma, iNKT cells active crosstalk with TANs skews their cytokine production from tolerogenic to pro-inflammatory ^23^. In inflammation, neutrophils regulate iNKT cell extravasation in the lung parenchyma ^25^ and license them to limit autoimmune responses in the spleen ^24^. In our CRC mouse models, the lack of iNKT cells significantly reduced the abundance of TANs and deeply affected their phenotype. We showed that iNKT cells are necessary to imprint an activated PMN-MDSCs gene signature ^26^ with immune suppressive activity as demonstrated by functional assays. The finding that iNKT cells favor neutrophil trafficking within cancer lesions worsening tumor burden was however in sharp contrast with previous studies showing that neutrophil infiltration is associated with a better survival in CRC ^31–33^. Nonetheless, the beneficial role of neutrophils in CRC was dependent on iNKT cells, since we found that the concomitant high infiltration of iNKT cells was a negative prognostic factor in our patients’ cohort and in the TCGA database. These findings parallel our *in vivo* experiments where the treatment of tumor-bearing mice with αGalCer reduced tumor burden, suggesting that modulating iNKT cell activation status may be considered a valid therapeutic option to restore their cytotoxic and anti-tumoral functions.

Our study showed that tumor infiltrating iNKT cells can first contribute to the remodeling of the TME by recruiting TANs, thereby sculpting the cancer developmental trajectory. Our findings uncover cellular and molecular mechanisms through which the iNKT-TAN axis can suppress antitumor immunity in CRC (Figure 6) and support the targeted manipulation of iNKT cells’ function to improve cancer immunotherapies such as checkpoint inhibitors and CAR/TCR-engineered T/iNKT cells ^41^.

**Figure 6:**
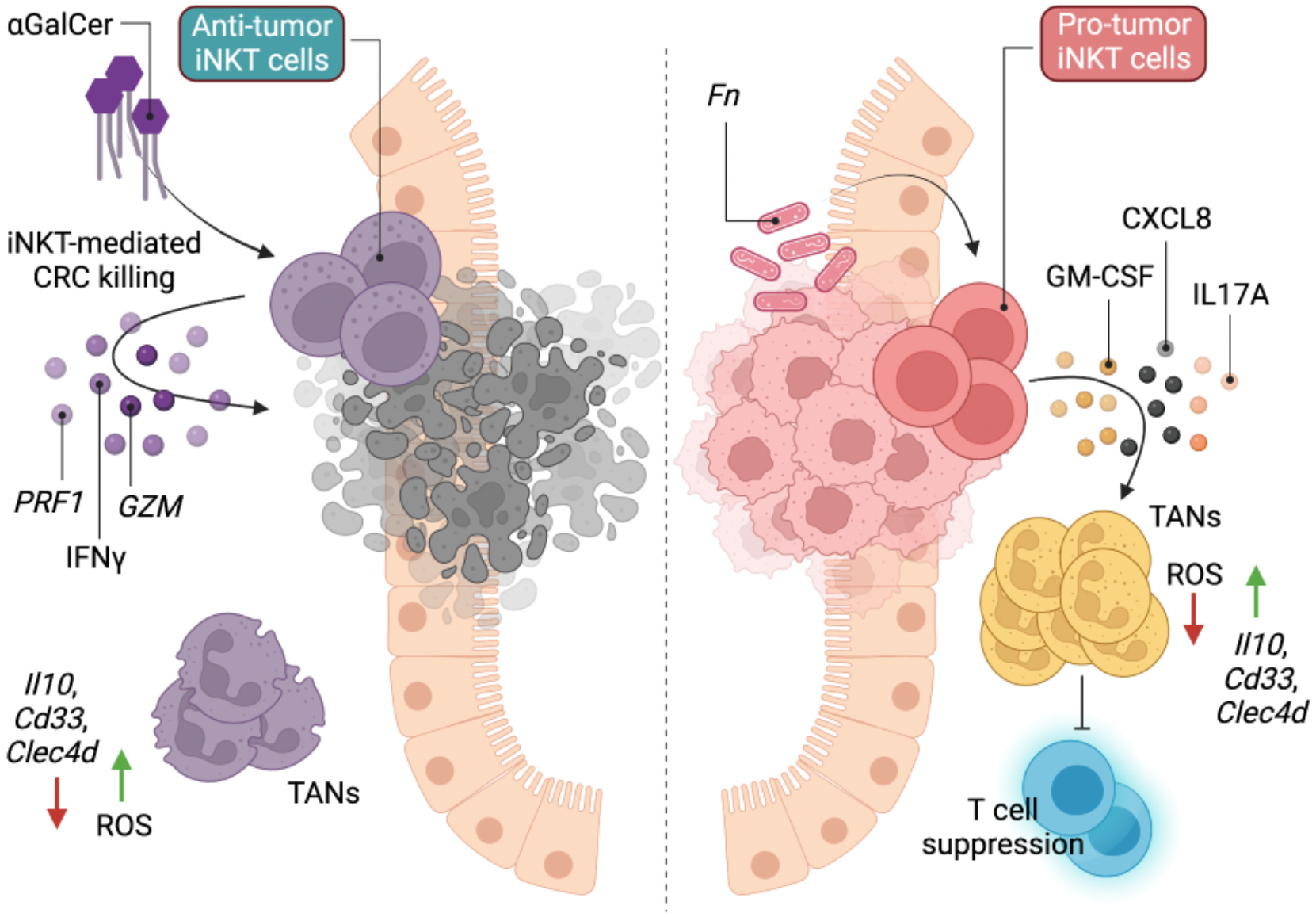
Proposed model for iNKT cell-mediated pro/antitumor immunity in CRC. The CRC-associated pathobiont *F. nucleatum* (*Fn*) impairs iNKT cell cytotoxic functions and promotes a pro-inflammatory phenotype in iNKT cells. Moreover, iNKT cell conditioning by *F. nucleatum* promotes iNKT cell-mediated recruitment of neutrophils with phenotypic and functional characteristics ascribable to polymorphonuclear myeloid-derived suppressor cells (PMN-MDSCs) in the TME (panel on the right of the dotted line). Our findings indicate that restoring the cytotoxic potential of iNKT cells by treating them with αGalCer leads to control of tumor growth (panel on the left of the dotted line).

## Materials & Methods

### Human Samples

Tumors and adjacent non-tumoral colon tissues were collected with informed consent from patients (n = 118) diagnosed with colorectal adenocarcinoma between January 2017 and July 2022 undergoing surgical resection at IRCCS Policlinico Ospedale Maggiore, Milan, Italy, as approved by the Institutional Review Board (Milan, Area B) with permission number 566_2015. AJCC IV patients ^42^ have been excluded from this study. Patient clinical data are summarized in Table 1.

### Human cells isolation

Tumoral samples were taken transversally to collect both marginal and core tumor zone. Normal adjacent tissues were sampled at least 10 cm from the tumor margin toward the ileum. Human lamina propria mononuclear cells (LPMCs) were isolated as previously described ^43^. Briefly, the dissected intestinal mucosa was freed of mucus and epithelial cells in sequential steps with DTT (0.1 mmol/l) and EDTA (1 mmol/l) (Sigma-Aldrich) and then digested with collagenase D (400 U/ml) (Worthington Biochemical Corporation) for 5 h at 37°C in agitation. LPMCs were then separated with a Percoll gradient.

### Neutrophil isolation

Neutrophils were isolated from whole blood samples by dextran sedimentation (4% diluted in HBSS) ^44^. Red blood cells were lysed using ACK lysis buffer (Life Technologies) and neutrophils separated with Percoll gradient.

### Generation of iNKT cell lines

Human iNKT cell lines were generated from sorted CD45^+^CD3^+^CD1d:PBS57Tet^+^ cells isolated from total LPMCs or PBMCs, as previously described ^7^. Sorted iNKT cells were stimulated with phytohemagglutinin (PHA, 1 μg·mL-1, Sigma-Aldrich) and irradiated peripheral blood feeders in a 2:1 iNKT : feeder ratio. PBMCs used as feeders were irradiated at 12.5 Gy. Stimulated cells were then expanded for 15 days by subculturing them every 2–3 days and maintained in RPMI-1640 medium with stable glutamine, 5% v/v human serum, and 100 IU·mL-1 IL-2 (Proleukin).

### Mice

6 to 8 weeks old C57BL/6 mice (Charles River, IT) were housed at the IEO animal facility in SPF conditions. *Traj18*^-/-^ and *CD1d*^-/-^ mice were bred and maintained at the IEO animal facility in SPF conditions. Aged-matched male and female mice were used for experiments. Animal experimentation was approved by the Italian Ministry of Health (Auth. 10/21 and Auth. 1217/20) and by the animal welfare committee (OPBA) of the European Institute of Oncology (IEO), Italy.

### Murine models of carcinogenesis

#### AOM-DSS model

7 weeks old mice were injected intraperitoneally with 10 mg/kg body weight Azoxymethane (AOM, Merck), dissolved in isotonic saline solution. After 7 days, mice were given 1% (w/v) dextran sodium sulfate (DSS MW 40 kD; TdB Consultancy) in their drinking water for 7 days followed by 14 days of recovery. The cycles were repeated 2 or 3 times and mice sacrificed at day 49 or 70.

#### MC38 model

7 weeks old mice were injected subcutaneously with 4 × 10^5^ MC38 cells. Tumor volume (V) was calculated from the caliper measurements using the following formula: V = (W^2^ × L)/2, where W is the tumor width and L is the tumor length ^45^. 2μg of αGalCer was injected intraperitoneally at day 0, 3 and 6. iNKT reconstitution in *Traj18*^-/-^ mice was performed co-inoculating freshly sorted splenic iNKT with MC38 cells at 1:4 ratio. Tumor-bearing animals were sacrificed after 15 days or earlier when showing any sign of discomfort.

### Murine colonoscopy

Colonoscopy was performed weekly for tumor monitoring using the Coloview system (TP100 Karl Storz, Germany). During the endoscopic procedure mice were anesthetized with 3% isoflurane.

### Murine cells isolation

Single-cell suspensions were prepared from the colon of C57BL/6, *Traj18*^-/-^ and *CD1d*^-/-^ mice as previously described ^9^. Briefly, cells were isolated via incubation with 5 mM EDTA at 37°C for 30 min, followed by mechanical disruption with GentleMACS (Miltenyi Biotec). After filtration with 100-μm and 70-μm nylon strainers (BD), the LPMC were counted and stained for immunophenotyping. MC38 tumors were digested with 1.5 mg/mL collagenase in RPMI-1640+10%FBS at 37°C for 60min. Cell suspension was filtered through 70-μm cell strainers, washed, counted and stained for multiparametric flow cytometry.

### *Fusobacterium nucleatum* culture condition

*F. nucleatum* strain ATCC25586 was maintained on Columbia agar supplemented with 5% sheep blood or in Columbia broth (Difco, Detroit, MI, USA) under anaerobic conditions at 37 °C. Columbia broth was supplemented with hemin at 5 μg·mL-1 and menadione at 1 μg·mL-1. Bacterial cell density was adjusted to 1 × 10^7^ CFU·mL-1 and heat-killed at 95°C for 15min before being stored at −80 °C until use in downstream experimentation.

### αGalCer and Fn-priming of iNKT cell

Monocyte derived dendritic cells (moDCs) were pulsed with αGalCer (100ng/ml) or with heat-inactivated *Fusobacterium nucleatum* (*Fn*) (4 × 10^5^ CFU) and co-cultured with iNKT cells (2 × 10^5^ cells) in a 2:1:4 iNKT:moDC:*Fn* ratio in RPMI-1640 supplemented with 10%FBS, Pen/Strep. After 24 h, iNKT cell activation status was estimated by intracellular staining.

### iNKT cell cytotoxicity assay

iNKT cell cytotoxicity toward the human CRC cell lines Colo205 and RKO (American Type Culture Collection, ATCC) was performed as previously described ^17^. Cytotoxicity was assessed using the Cytotoxicity Lactate Dehydrogenase (LDH) Assay Kit-WST (nonhomogeneous assay, Dojindo, EU) following the manufacturer’s instructions. All experimental conditions were performed in duplicate. Cancer cells (2.5 × 10^4^ cells/well) were incubated at 37 °C for 4 h with effector cells at effector:target ratio of 8:1. Supernatants were collected and plated in optically clear, 96-well plates, and absorbance at 490 nm was measured using a GloMax Microplate Reader (Promega, Madison, WI, USA) after the colorimetric reaction for LDH detection was finished. The percentage of cytotoxicity was calculated as follows: (test well – spontaneous release control)/(maximal release control – spontaneous release control) × 100.

### iNKT-Neutrophil co-culture assay

αGalCer or *F. nucleatum* primed-iNKT cells (2 × 10^5^ cells) were co-cultured with freshly isolated neutrophils in a 1:1 ratio, in RPMI-1640 supplemented with 10% FBS. After 24h cells were stained for extracellular markers expression and ROS detection.

### In-vitro suppression assay

Naïve CD4^+^T cells were isolated from PBMCs of healthy donors (CD4 näive human microbeads, Miltenyi Biotech). Cells were labelled with 1 nM Far Red CellTrace (ThermoFisher), re-suspended in medium containing hIL2 (Proleukin) and anti-CD28 antibody (2 μg/ml, Tonbo) and plated in 96-well plates (NUNC Maxisorp) pre-coated with anti-CD3 antibody (2 μg/ml, Tonbo) at a concentration of 2.5 × 10^4^ cells/well. Freshly isolated neutrophils were co-cultured with T cells at a 1:1 ratio and the culture supernatant from NS, αGalCer or *Fn*-primed iNKT cells were added at a final concentration of 10%. After 5 days, proliferating naïve CD4^+^T cells were labelled with Zombie vital dye (Biolegend) and analyzed with a BD FACS Celesta. The suppression index was calculated using the FlowJo Proliferation Modeling tool and normalized on minimum proliferation levels.

### Neutrophil migration assay

Freshly isolated neutrophils were first pre-incubated 20 min at 37 °C with Reparixin (20 μM) or RPMI-1640+2%FBS and then seeded on top of a 3μm-pore transwell (SARSTEDT) in 200μl of RPMI-1640+2%FBS. 500μl of chemoattracting medium *i.e*., the culture supernatant of activated iNKT cell lines diluted 10% in RPMI-1640+2% FBS (see the αGalCer and *Fn*-priming of iNKT cell protocol) was added on the bottom of the transwell and neutrophils’ migration was allowed for 4 hours at 37°C. RPMI-1640+10% FBS was used as positive control. After 4 hours of incubation, the total number of cells on the bottom of the plate were stained and counted using the FACSCelesta flow cytometer (BD Biosciences, Franklin Lakes NJ, USA) with plate-acquisition mode and defined volumes.

### Neutrophil survival assay

Freshly isolated neutrophils were cultured with RPMI-1640+10% FBS supplemented with the culture supernatants (10%) from αGalCer or *Fn*-primed iNKT cells for 16h at 37°C. Cells were then stained with FITC Annexin V Apoptosis Detection Kit with 7-AAD (Biolegend) following manufacturer’s instruction and acquired at a FACS Celesta flow cytometer (BD Biosciences, Franklin Lakes NJ, USA).

### Respiratory Burst Assay

ROS production was quantified using the Neutrophil/Monocyte Respiratory Burst assay (Cayman) following manufacturer’s instructions.

### ELISA assay

Detection of IL8/CXCL8 in culture supernatants was performed using the OptEIA Human IL-8 kit (BD Biosciences), according to manufacturer’s instruction.

### Gut microbiota-priming of murine iNKT cells

Splenic iNKT cells were isolated from C57BL/6 mice sorting CD45^+^CD3^+^CD1d:PBS57Tet^+^ cells upon enrichment through B cells exclusion (Mouse CD19 microbeads, Miltenyi Biotec). Bone marrow derived dendritic cells (BMDCs) from C57BL/6 mice were pulsed with heat-inactivated fecal microbiota of controls or AOM-DSS treated C57BL/6, *Traj18*^-/-^ and *CD1d*^-/-^ mice and co-cultured with freshly isolated splenic iNKT cells (2 × 10^5^ cells) in a 2:1:10 iNKT:BMDC:microbiota ratio in RPMI-1640 supplemented with 10% FBS and Pen/Strep solution. After 24h, iNKT cell activation status was estimated by intracellular staining. Fecal samples were resuspended 1:10 (w/v) in PBS and filtered through a 0.75 μm filter to remove large debris; microbiota cell density was quantified by qPCR ^46^, adjusted to 2 × 10^7^ CFU·mL-1 and heat-killed at 95°C for 15min before being stored at −80 °C until use in downstream experimentation.

### Flow Cytometry

iNKT cells were stained and identified using human or mouse CD1d:PBS57 Tetramer (NIH Tetramer core facility) diluted in PBS with 1% heat-inactivated FBS for 30 min at 4°C. For intracellular cytokine labeling cells were incubated for 3 h at 37°C in RPMI-1640+10% FBS with PMA (10μg/ml, Merck), Ionomycin (1μg/ml, Merck) and Brefeldin A (10 μg/ml, Merck). Before intracellular staining cells were fixed and permeabilized using Cytofix/Cytoperm (BD). Samples were analyzed with a FACSCelesta flow cytometer (BD Biosciences, Franklin Lakes NJ, USA). Data were analyzed using the FlowJo software (Version 10.8, TreeStar, Ashland, OR, USA). For the multi-dimensional analysis using t-SNE visualization and Phenograph clustering refer to the dedicated section in methods.

### Multi-dimensional flow cytometry analysis

FCS files were upload in FlowJo software (Version 10.8) and data were compensated manually according to the software usage. One fluorescence parameter per laser was then queried for irregularities during the acquisition by checking it over the time parameter. If variations in the flow stream were detected, they were excluded from the analysis. Data were cleaned for antibodies aggregates by checking each parameter in a bimodal plot. Gate on singlets, on viable lymphocytes and subsequently on CD3^+^cells were applied. CD3^+^ populations were down-sampled to 5000 events per sample using the DownSample plugin (Version 3.3.1) of FlowJo to create uniform population sizes. Down-sampled populations were exported as FCS files with applied compensation correction. Files were then uploaded to RStudio environment (Version 3.5.3) using the flowCore package (Version 1.38.2). Data were transformed using logicleTransform() function present in the flowCore package. To equalize the contribution of each marker they were interrogated for their density distribution using the densityplot() function of the flowViz package (Version 1.36.2). Each marker was normalized using the Per-channel normalization based on landmark registration using the gaussNorm() function present in the package flowStats (Version 3.30.0). Peak.density, peak.distance and number of peaks were chosen according to each marker expression. Normalized files were analyzed using the cytofkit package through the cytofkit_GUI interface. For data visualization we used the t-Distributed Stochastic Neighbor Embedding (t-SNE) method, while for clustering we used the Phenograph algorithm. t-SNE plots were visualized on the cytofkitShinyAPP with the following parameters: perplexity=50, iterations=1000, seed=42, k=50. FCS for each cluster were generated and re-imported in FlowJo to be manually analyzed for the determination of the integrated MFI. The iMFI of different markers was scaled from 0 to 1 and used to identify Phenograph clusters ^47^.

### 16S rRNA gene sequencing and data analysis

Intestinal mucus scraped from the human and murine colons were stored at −80°C until DNA extraction. DNA extraction, 16S rRNA gene amplification, purification, library preparation and pair-end sequencing on the Illumina MiSeq platform were performed as previously described ^9^. Reads were pre-processed using the MICCA pipeline (v.1.7.0) (http://www.micca.org) ^48^ as previously described ^9^. Differential abundance testing was carried out using the R package DESeq2 ^49^ using the non-rarefied data ^50^. P-values were False Discovery Rate corrected using the Benjamini-Hochberg procedure implemented in DESeq2.

### Bulk RNA sequencing of human iNKT cells

Total RNA (from 1 × 10^6^ cells) was isolated with the RNeasy kit (Qiagen) and RNA quality was checked with the Agilent 2100 Bioanalyzer (Agilent Technologies). 0.5-1 μg were used to prepare libraries for RNA-seq with the Illumina TruSeq RNA Library Prep Kit v2 following the manufacturer’s instructions. RNA-seq libraries were then run on the Agilent 2100 Bioanalyzer (Agilent Technologies) for quantification and quality control and pair-end sequenced on the Illumina NovaSeq platform.

### Bulk RNA sequencing of sorted neutrophils

Total RNA from ~ 5 × 10^5^ neutrophils (CD45^+^Lin^-^CD11b^+^Ly6G^+^) was isolated with the RNeasy micro kit (Qiagen) and RNA quality was checked with the Agilent 2100 Bioanalyzer (Agilent Technologies). Sequencing libraries were prepared by using the NEBNext® rRNA Depletion Kit v2 and the NEBNext® Ultra™ II Directional RNA Library Prep kits following manufacturer’s instructions. RNA-seq libraries were then run on the Agilent 2100 Bioanalyzer (Agilent Technologies) for quantification and quality control and pair-end sequenced on the Illumina NovaSeq platform.

### RNA sequencing data analysis

RNA-seq reads were preprocessed using the FASTX-Toolkit tools. Quality control was performed using FastQC. Pipelines for primary analysis (filtering and alignment to the reference genome of the raw reads) and secondary analysis (expression quantification, differential gene expression) have been integrated and run in the HTS-flow system ^51^. Differentially expressed genes were identified using the Bioconductor Deseq2 package ^49^. P-values were False Discovery Rate corrected using the Benjamini-Hochberg procedure implemented in DESeq2. Functional enrichment analyses to determine Gene Ontology categories and KEGG pathways were performed using the DAVID Bioinformatics Resources (DAVID Knowledgebase v2022q2) (https://david.ncifcrf.gov) ^52,53^.

### Single cell RNA sequencing data analysis

Raw count matrices were downloaded from the Gene Expression Omnibus (GEO): accession number GSE163834. PMNs from spleen of naïve mice (n=3), spleen of tumour-bearing mice (n=3), and tumours (n=3) were analysed, yielding to a total of 66,854 single cells. Data processing and analysis was performed using the Seurat workflow ^54^. Counts were normalized and log-transformed using sctransform ^55^, while regressing out UMI counts and the percentage of mitochondrial counts. Highly variable genes were used to perform principal component analysis (PCA) and principal components (PCs) covering the highest variance in the dataset were selected. PCs were fed to Harmony ^56^ for batch correction. Clusters were identified using the shared nearest neighbour (SNN) modularity optimization-based clustering algorithm, followed by Louvain community detection. 14 clusters were identified. Two of these clusters were characterized as B cells and macrophages (*Cd79a, Cd79b* for B cells, and *Csf1r, Mafb, Adgre1* for macrophages) and were removed from the analysis. All the others clusters were annotated into PMN1, PMN2, and PMN3, based on the expression levels of the markers described in ^26^. The AddModuleScore function was used to calculate the module score for the two signatures of interest derived from bulk RNA-seq data. The signatures included the genes upregulated in C57BL/6 mice (FDR p < 0.1 e log_2_FC < −1) and the genes upregulated in *Traj18*^-/-^ mice (FDR p < 0.1 e log_2_FC > 1).

### Statistical analysis

Statistical tests were conducted using Prism (Version 8.2.0, GraphPad) software or the R software (version 3.6.2). Paired, non-parametric Wilcoxon test was used to compare non-tumoral and tumoral tissues, both in human and murine samples. The Mann-Whitney U test was used for unpaired comparisons. Spearman’s correlation coefficient was used for the analysis of correlations. Random Forest ^57^ analysis of flow cytometric data from innate immune cells was performed using the randomForest R package; permutation tests with 1000 permutations were performed to assess model significance ^58^. Kaplan-Meier analysis were carried out using the R packages *survival* (version 3-2-11) and *survminer* (version 0.4.9). Statistical analyses were always performed as two-tailed. P-values were corrected for multiple comparisons and considered statistically significant with p < 0.05. ***p < 0.001; **p < 0.01; *p < 0.05.

## Acknowledgments

We thank the IEO Animal Facility for the excellent animal husbandry, the IEO Genomic Unit for supporting in high throughput sequencing, the NIH Tetramer Facility for providing human and murine CD1d:PBS57 tetramers. We are grateful to the équipe of the General and Emergency Surgery Unit, Ospedale Maggiore Policlinico, Milano for their tireless work. We thank Dr. Paolo Dellabona for providing *CD1d*^-/-^ and *Traj18*^-/-^ mice. We thank Prof. Maria Rescigno, Prof. Massimo C. Fantini, Dr. Matteo Marzi and Dr. Roberto Gianbruno for the helpful discussions and support in the set-up of sequencing experiments. We thank Claudia Burrello and Erika Mileti for initial set-ups of the experiments. Schemes in Fig. 2B and 4A were created using icons from the Noun Project (https://thenounproject.com/). Figure 6 was created with BioRender.com (https://biorender.com/).

## Supplementary Material

**Fig. S1:**
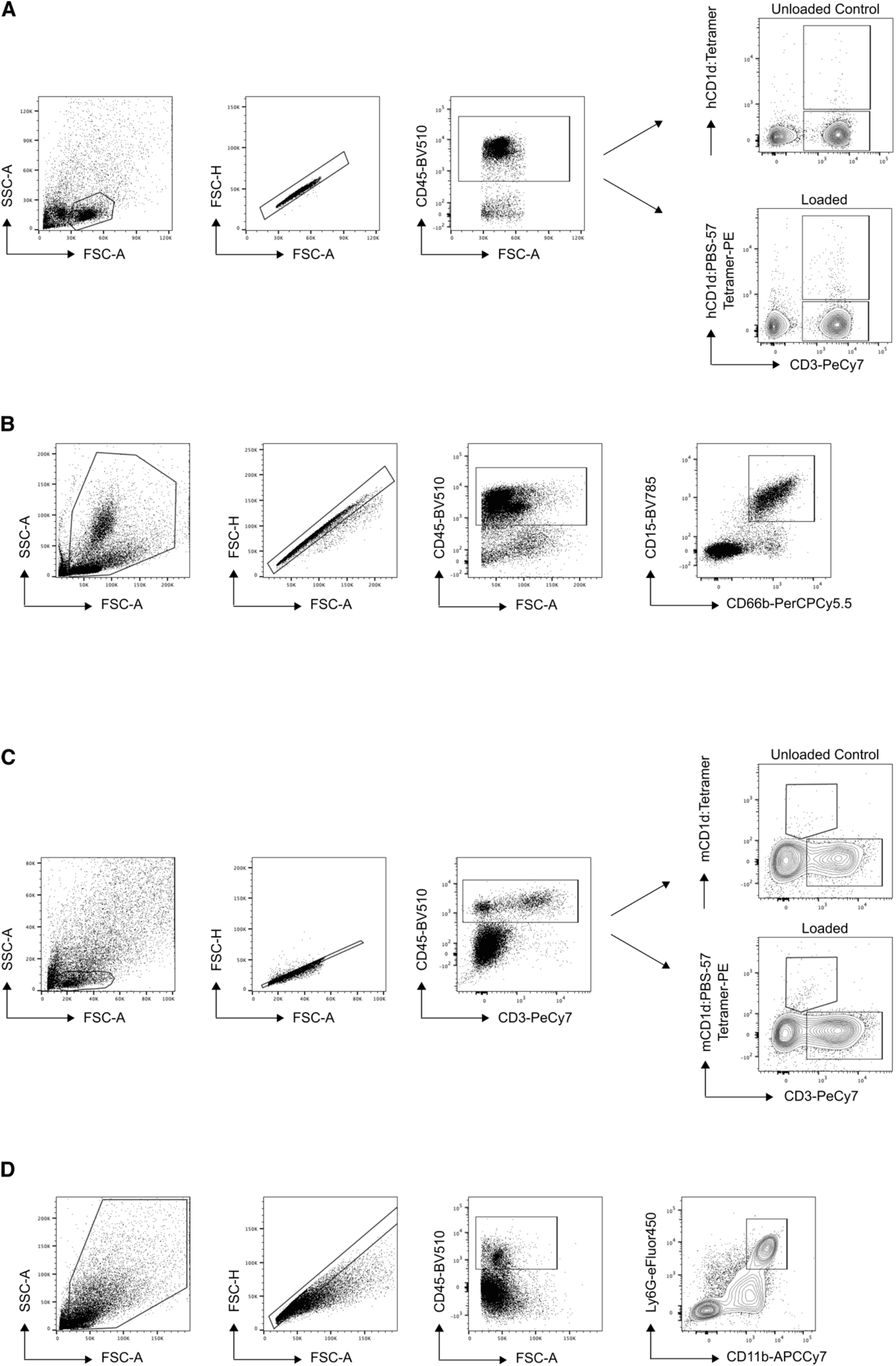
Gating strategies for iNKT cells and neutrophils. **(A)** iNKT cells and **(B)** neutrophils gating strategies for specimens of CRC patients. **(C)** iNKT cells and **(D)** neutrophils gating strategies for murine samples. Unloaded CD1d tetramers served as control.

**Fig. S2:**
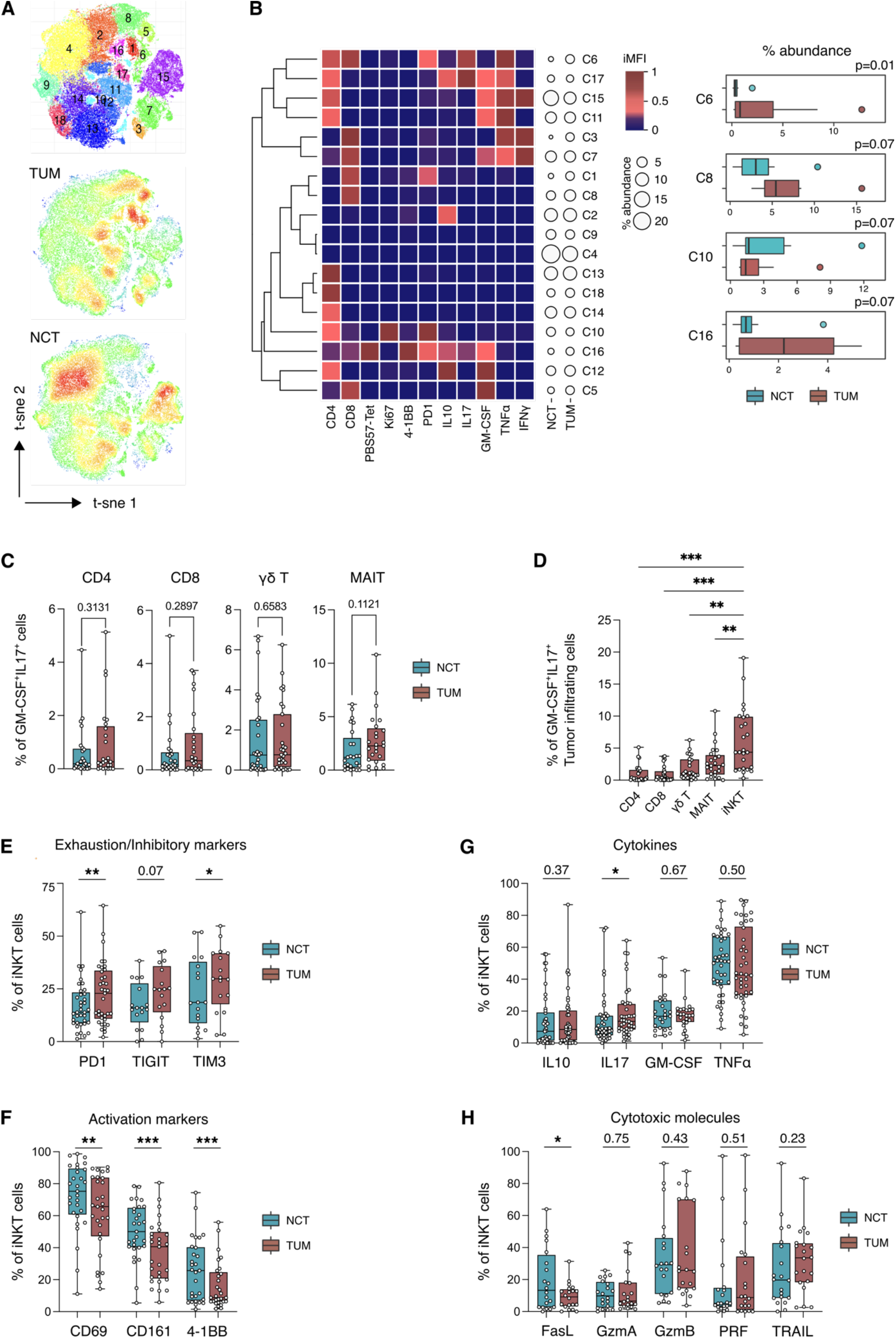
Immunophenotyping of iNKT cells in NCT and TUM. **(A)** t-SNE map of CD3^+^T cells based on Phenograph clustering in NCT and TUM. **(B)** Heatmap of scaled integrated MFI data from Phenograph clustering analysis; relative abundance of identified clusters from NCT and TUM is also shown. **(C)** Frequency of IL17^+^GM-CSF^+^ conventional (CD4^+^ and CD8^+^) and unconventional (γδ and MAIT) T cells infiltrating NCT and TUM (n=25). **(D)** Frequency of tumor infiltrating IL17^+^GM-CSF^+^ conventional (CD4^+^ and CD8^+^) and unconventional (γδ, MAIT and iNKT) T cells (n=25). **(E)** Frequency of PD1^+^, TIGIT^+^ and TIM3^+^ iNKT cells infiltrating NCT and TUM (n=16-37). **(F)** Frequency of CD69^+^, CD161^+^ and CD137^+^ iNKT cells infiltrating NCT and TUM (n=29) **(G)** Frequency of IL10^+^, IL17^+^, GM-CSF^+^ and TNFα^+^ iNKT cells infiltrating NCT and TUM (n=42). **(H)** Frequency of FasL^+^, GzmA^+^, GzmB^+^, Perforin^+^ (PRF) and TRAIL^+^ iNKT cells infiltrating NCT and TUM (n=20). P < 0.05 (*), P < 0.01 (**), P < 0.001(***); Wilcoxon signed rank test and Friedman test.

**Fig. S3:**
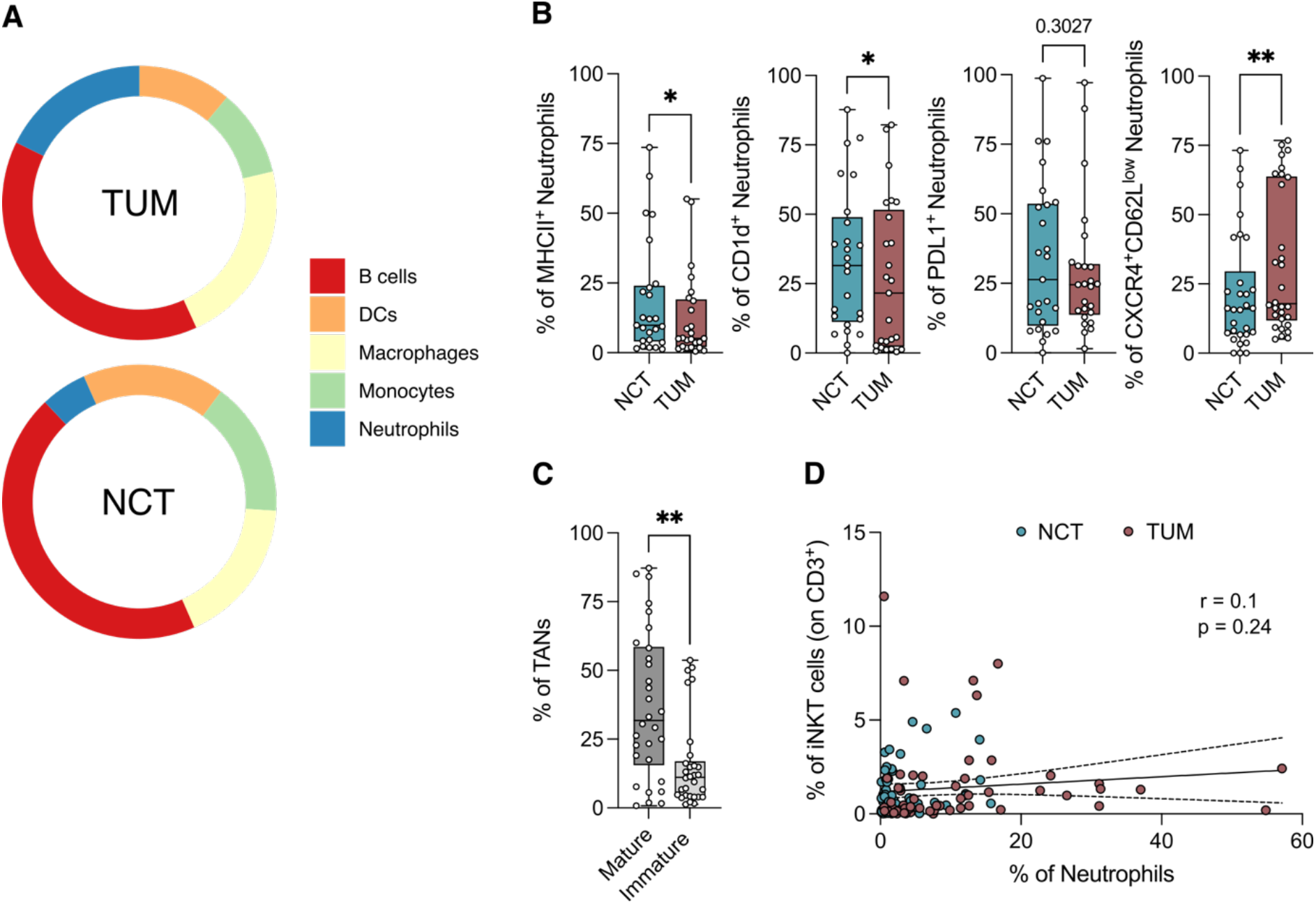
Neutrophil characterization in NCT and TUM. **(A)** Pie chart of myeloid and B cells frequency in NCT and TUM samples (**B)** Frequency of MHCII^+^, CD1d^+^, PD-L1^+^ and CXCR4^+^CD62L^low^ neutrophils in NCT and TUM (n=25-30). (**C)** Frequency of mature (CD33^med^CD10^+^CD16^+^) and immature (CD33^med^CD10^+^CD16^-^) TANs (n=30). (**D)** Spearman’s correlation analysis of tissue infiltrating iNKT cells and neutrophils (n=58). P < 0.05 (*), P < 0.01 (**); Wilcoxon signed rank and Mann-Whitney tests.

**Fig. S4:**
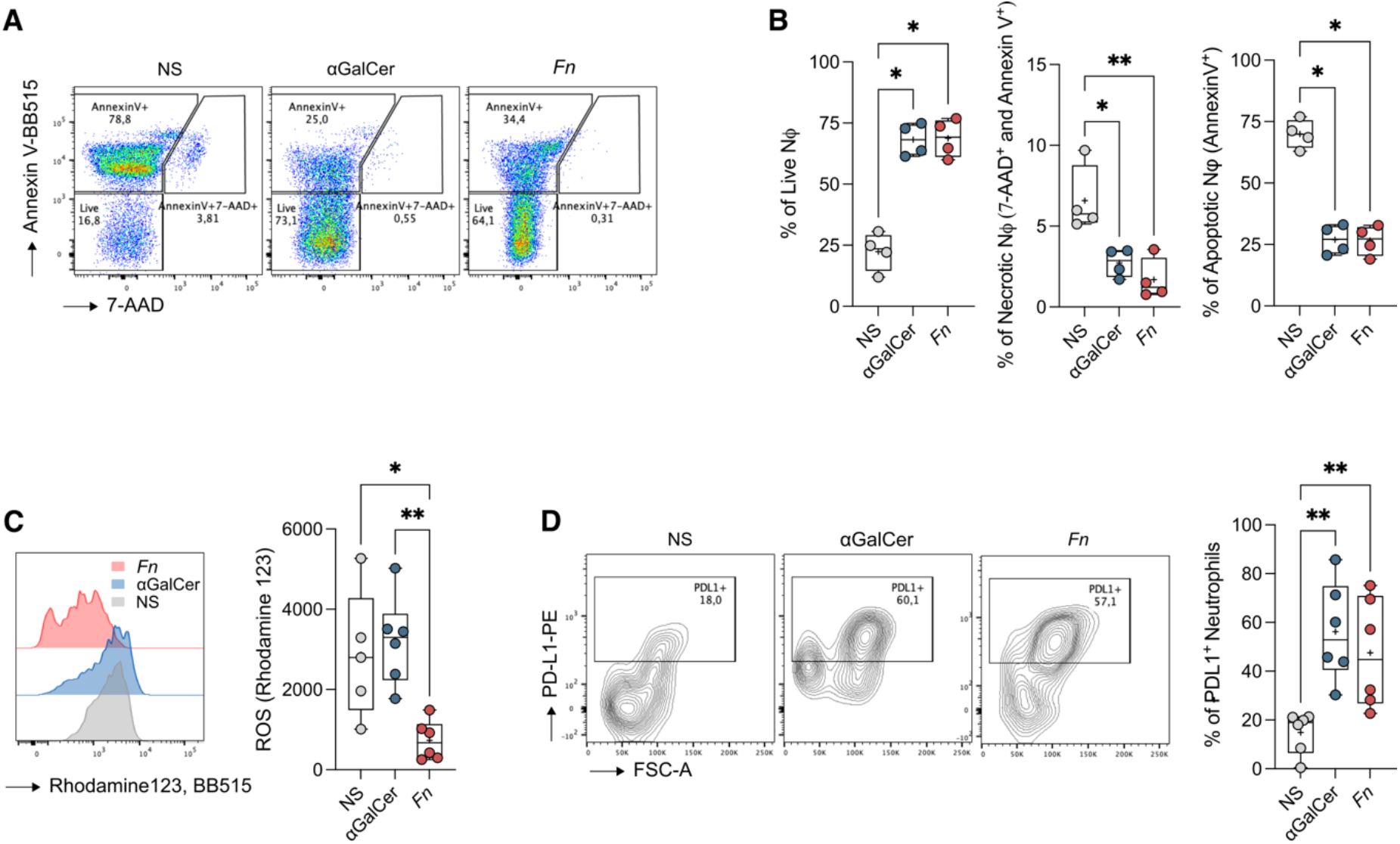
iNKT cell-primed supernatant affects neutrophil survival and function. **(A)** Representative plots of Annexin V and 7-AAD staining from neutrophils exposed to the culture supernatants of unstimulated (NS), αGalCer or *F. nucleatum* (*Fn*) primed-iNKT cells. (**B)** Frequency of live (7-AAD^-^, Annexin V^-^), necrotic (7-AAD^+^ and AnnexinV^+^) and apoptotic (7-AAD^-^, Annexin V^+^) neutrophils. (**C)** Respiratory burst assay quantification and (**D)** frequency of PD-L1^+^ cells from neutrophils exposed to the culture supernatants of unstimulated (NS), αGalCer or *F. nucleatum* (*Fn*) primed-iNKT cells with representative plots. P < 0.05 (*), P < 0.01 (**); Kruskal-Wallis test. Data are representative of at least three independent experiments.

**Fig. S5:**
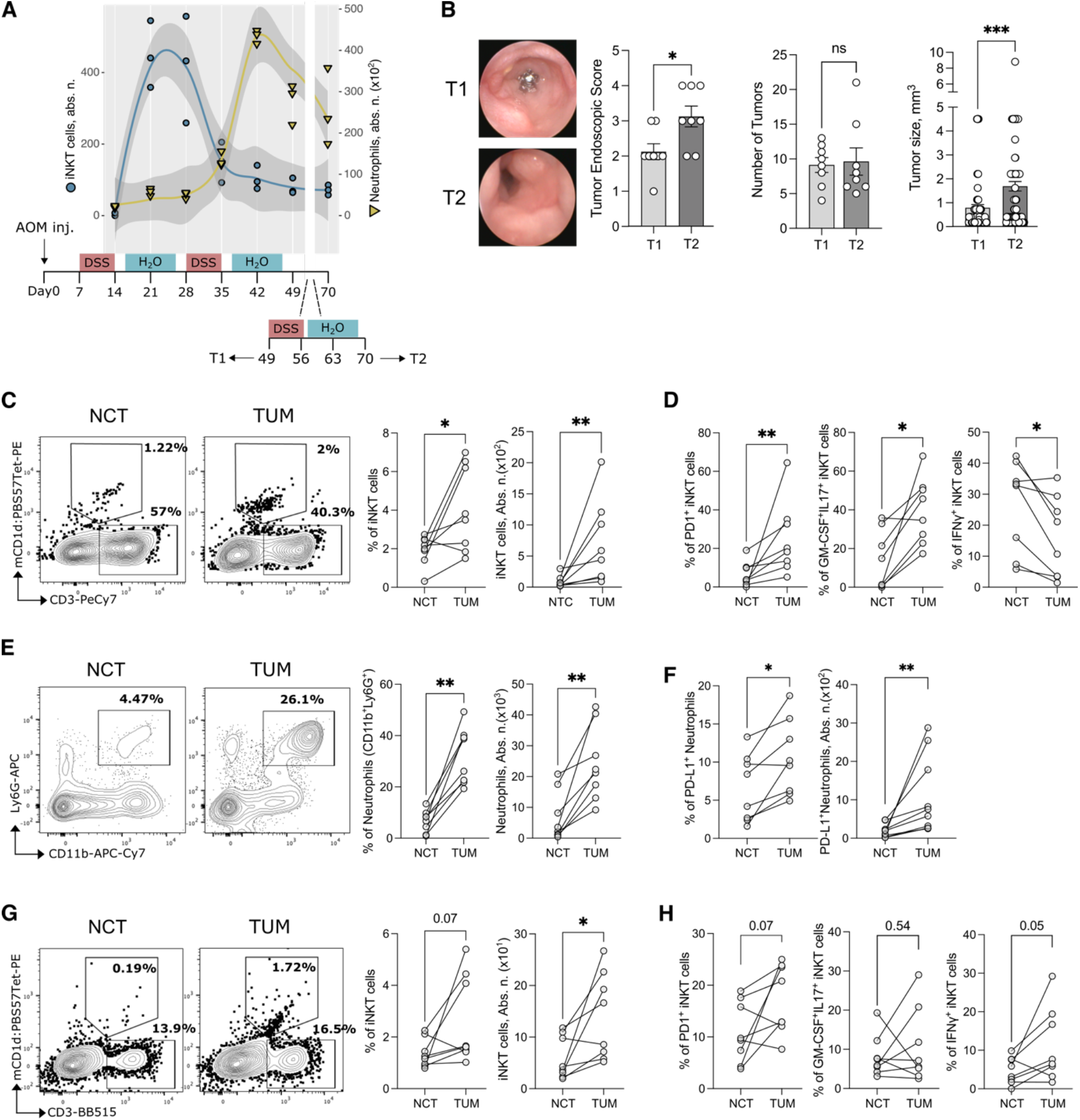
Phenotypic evaluation of iNKT cells and neutrophils in AOM-DSS treated mice. **(A)** iNKT cells and neutrophils dynamic infiltration in tumors upon AOM-DSS treatments with timeline scheme of the experimental protocol. **(B)** Tumor endoscopic score with representative endoscopic pictures, tumor numbers and size in AOM-DSS-treated mice at T1 and T2. **(C)** Frequency and absolute numbers of iNKT cells infiltrating NCT and TUM in AOM-DSS-treated mice at T1, with representative dot plots. **(D)** Frequency of PD-1^+^, GM-CSF^+^ and IFNγ^+^iNKT cells infiltrating NCT and TUM at T1. **(E)** Frequency and absolute numbers of neutrophils infiltrating NCT and TUM at T1, with representative plots. **(F)** Frequency and absolute numbers of PD-L1^+^ neutrophils infiltrating NCT and TUM at T1. **(G)** Frequency and absolute numbers of iNKT cells infiltrating NCT and TUM in the AOM-DSS model at T2, with representative dot plots. **(H)** Frequency of PD-1^+^, GM-CSF^+^ and IFNγ^+^iNKT cells infiltrating NCT and TUM in AOM-DSS-treated mice at T2. P < 0.05 (*), P < 0.01 (**), P < 0.001(***); Mann-Whitney test.

**Fig. S6:**
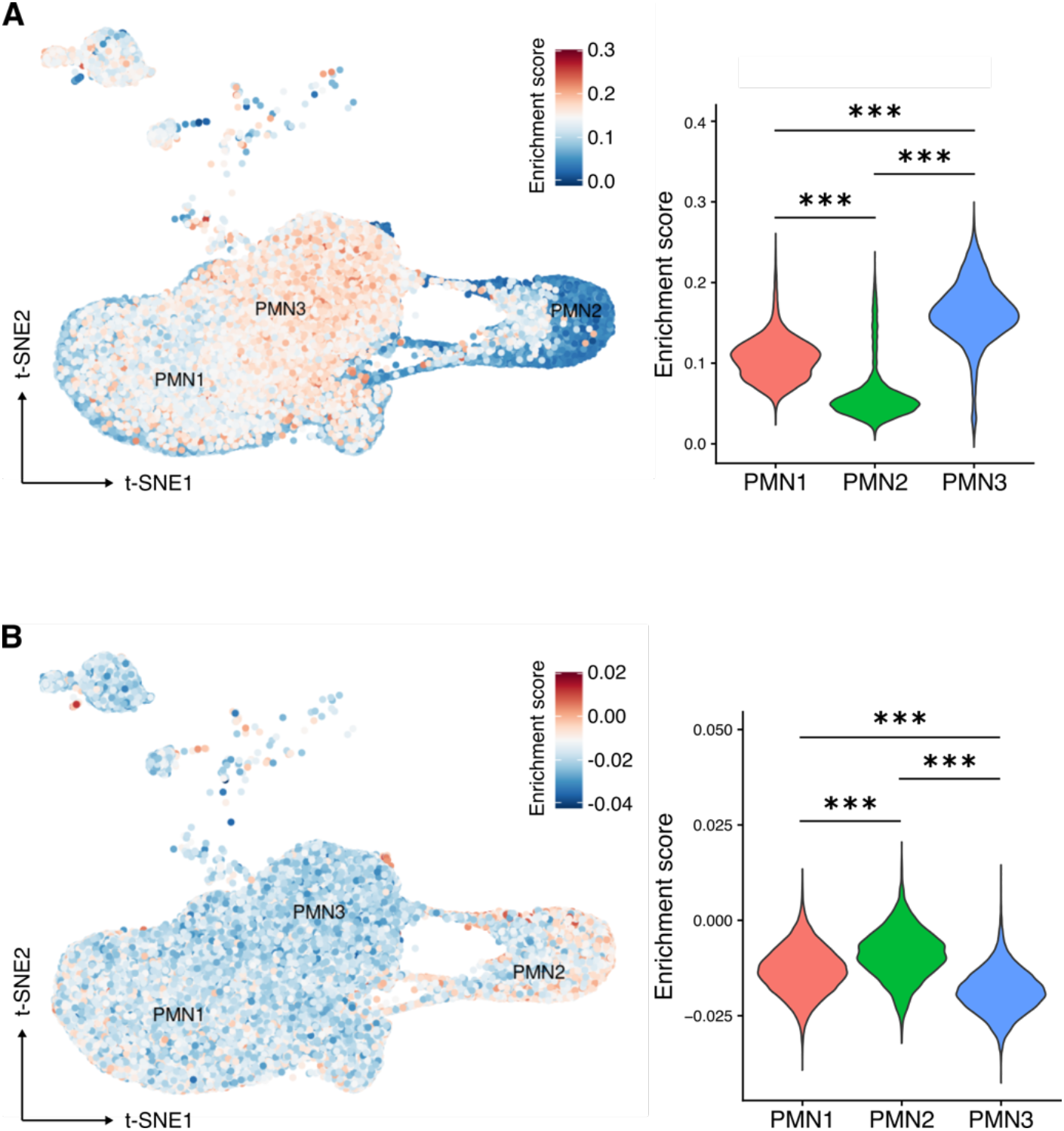
C57BL/6 and *Traj18*^-/-^ TANs gene signature associate with different populations of PMN-MDSC. Single-cell signature correlation (t-SNE overlay) calculated using the log_2_FC of the **(A)** C57BL/6 and **(B)** *Traj18*^-/-^ TANs gene expression signatures (genes with an FDR-corrected p-value < 0.1 and log_2_FC > |1|) and the relative expression values of the single cell PMN-MDSC data set from Veglia *et al*., 2021. P < 0.001 (***); Kruskal-Wallis test.

**Fig S7:**
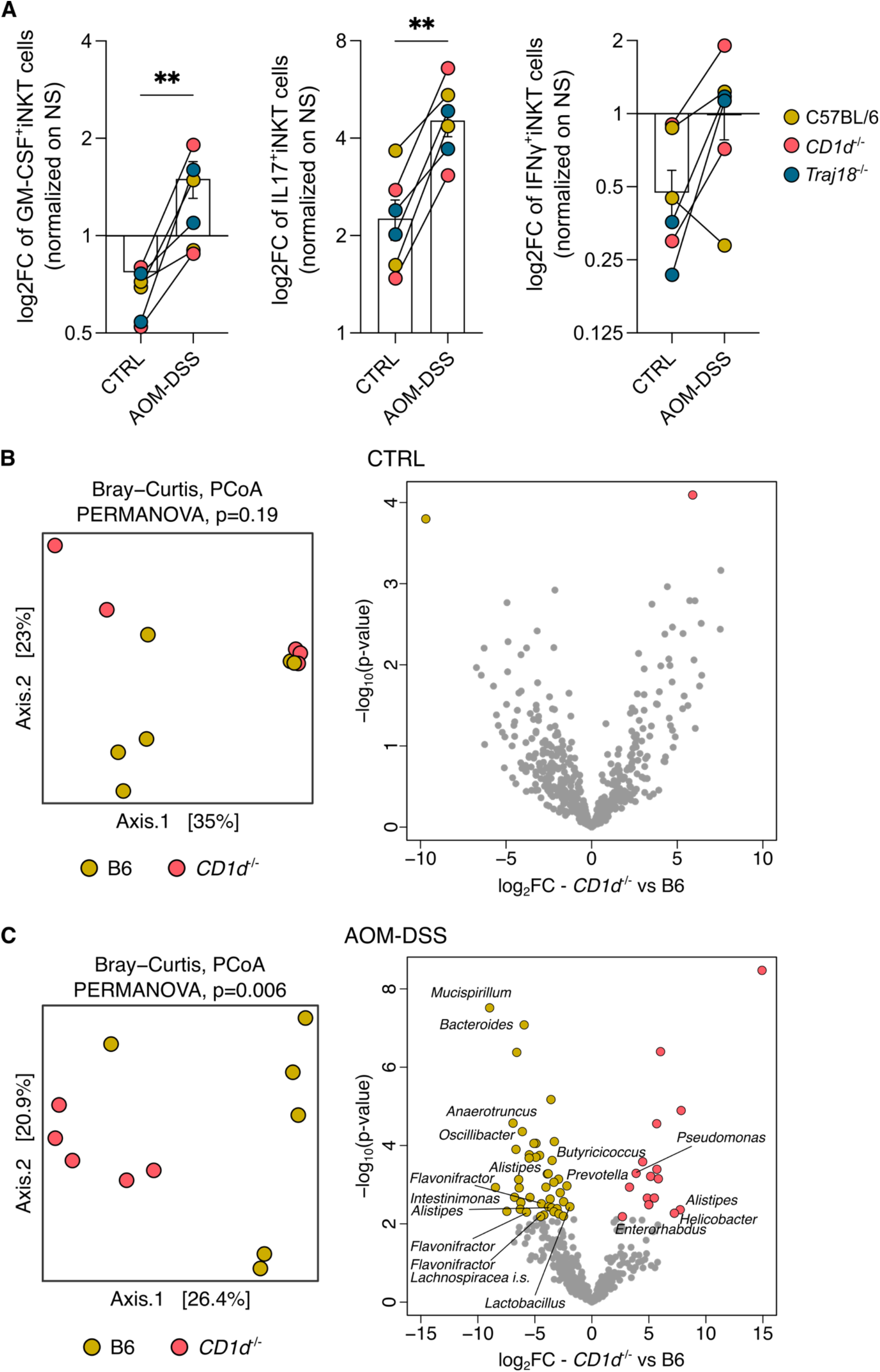
The CRC-associated microbiome promotes a GM-CSF/IL17 iNKT phenotype. **(A)** Frequency of GM-CSF^+^, IL17^+^ and IFNγ^+^iNKT cells sorted from spleen of healthy C57BL/6 mice upon BMDCs priming with the gut microbiota isolated from controls (CTRL) and AOM-DSS treated C57BL/6, *CD1d*^-/-^ and *Traj18*^-/-^ animals. Data shown from three independent experiments are expressed as log_2_fold-change (log_2_FC) normalized to unstimulated iNKT cell control. **(B-C)** PCoA of microbiota beta-diversity as measured by Bray-Curtis distance (left panels) and volcano plots (right panels) representing the significantly enriched bacterial taxa (FDR p <0.05) in **(B)** CTRL and **(C)** AOM-DSS treated C57BL/6 and iNKT-deficient *CD1d*^-/-^ mice. The names of the significantly enriched ASVs classified to the genus level are reported. P < 0.01 (**); Mann-Whitney tests.

